# Proportionality-based association metrics in count compositional data

**DOI:** 10.1101/2023.08.23.554468

**Authors:** Kevin McGregor, Nneka Okaeme, Reihane Khorasaniha, Simona Veniamin, Juan Jovel, Richard Miller, Ramsha Mahmood, Morag Graham, Christine Bonner, Charles N. Bernstein, Douglas L. Arnold, Amit Bar-Or, Janace Hart, Ruth Ann Marrie, Julia O’Mahony, E. Ann Yeh, Yinshan Zhao, Brenda Banwell, Emmanuelle Waubant, Natalie Knox, Gary Van Domselaar, Feng Zhu, Ali I. Mirza, Helen Tremlett, Heather Armstrong

## Abstract

**Motivation:** Compositional data comprise vectors that describe the constituent parts of a whole. Data arising from various -omics platforms such as 16S and RNA-sequencing are compositional in nature. However, correlations between features on raw counts have no meaningful interpretation. Metrics of proportionality were formulated to address this problem. However, there is an inherent bias that arises when calculating these metrics empirically on count-based measures due to variability in read depths.

**Results:** We quantify the bias introduced by empirically calculating proportionality-based association metrics in count data. Additionally, we propose a means of estimating these metrics within a logit-normal multinomial model in pursuit of more accurate estimates. The model-based estimates are shown to outperform empirical estimates in simulated data, and are additionally applied to a mouse embryonic stem-cell single-cell sequencing dataset as well as a pediatric-onset multiple sclerosis metagenomic dataset.

**Availability and Implementation:** An R package is available at https://CRAN.R-project.org/package=countprop.

**Supplementary information:** Supplementary data are available at Bioinformatics online.

## Introduction

Compositional data comprise vectors of quantitative measures that describe the constituent parts of a whole. The only information in compositional data is found in the *ratios* between the different parts of the vector; the individual measures are inherently uninterpretable [1]. Data arising from various omics platforms are compositional in nature— for instance, RNA-sequencing (RNA-Seq), 16S amplicon-sequencing, and single-cell RNA-sequencing are all examples of platforms that yield compositional data. The primary reason for the compositional classification of these platforms is that the number of sequences varies across samples due to technical artifacts in the sequencing process, meaning that within-sample counts must either be normalized to a unit sum, or log-ratio transformed in order to effect any kind of meaningful interpretation [9]. Some microbiome studies have relied on *rarefying* such data, which involves resampling the counts to a constant read depth across all samples. However, it has been recognized that such a procedure essentially amounts to discarding valid data and should therefore be discouraged [18].

This interpretability problem is particularly worrisome when considering associations between features—the prime example being Pearson’s correlation. Due to varying read depths, correlations on raw counts have no meaningful interpretation. Similarly, when normalizing the counts to proportions, a negative bias is introduced in the correlations as a result of the unit-sum constraint.

An appropriate class of measures based on *proportionality* has been established to address the issue of quantifying association in compositional data. The most basic such measure is the variation matrix **V** whose elements are defined as follows: suppose **p** = (*p*_1_, …, *p*_*D*_) is a random vector containing proportions such that 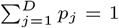. The element in row *j* and column *k* of **V**, which describes the association between features *j* and *k* is given by:

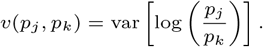

When *p*_*j*_ is exactly proportional to *p*_*k*_, we have that *v*(*p*_*j*_, *p*_*k*_) = 0. Hence, the smaller the value of *v*(*p*_*j*_, *p*_*k*_) the stronger the association between the abundances of features *j* and *k*. Note that the elements of **V** are non-negative, and thus the variation matrix does not give information about the direction of the associations of the pairs of features included in the matrix.

One important issue with the variation matrix is that its elements are variances of different pairs of features, meaning that they do not share a common scale. To address this issue, [7] developed alternative measures of proportionality.

The first is called *ϕ*, and is simply *v*(*p*_*j*_, *p*_*k*_) scaled by the variance of the logarithm of the first argument. It is defined as:

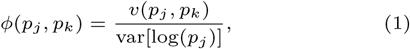

with *ϕ*(*p*_*j*_, *p*_*k*_) ≥ 0. The interpretation is the same as for *v*, only now it is scaled based on its first argument. The main downside of this metric is that it is not symmetric with regards to its arguments. A final proportionality metric is defined as:

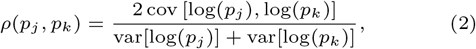

with −1 ≤ *ρ*(*p*_*j*_, *p*_*k*_) ≤ 1. If *ρ* ≈ 1, then *p*_*j*_ and *p*_*k*_ are strongly proportional, and if *ρ* ≈ −1, then *p*_*j*_ and *p*_*k*_ are strongly inversely proportional.

Metrics of proportionality recently gained more recognition, when [25] suggested their use to detect gene-gene associations in single-cell sequencing experiments. This publication compared numerous measures of association across multiple datasets by evaluating their ability to create functionally coherent single-cell gene co-expression networks. There was a clear advantage for the proportionality metrics across the datasets.

The one common property of these kinds of metrics is that they are *scale invariant*, which is the main property allowing their use in compositional data. However, classical techniques for the analysis of compositional data were developed with continuous measures in mind; for example, the constituent parts of a soil sample. Conversely, sequencing platforms provide data in the form of *counts*. A number of recent studies have shed light on problems that arise when applying traditional methods for compositional data analysis on count-based compositional data. [14] showed that proportionality cannot be exactly represented in count-based compositional data, which they referred to as *lattice* compositional data. [15] showed a similar phenomenon in logarithmic transformations in single-cell sequencing; varying read depths among cells led to systematic errors and spurious differences in expression. Relatedly, [6] presented novel modelling techniques in light of these issues in count data.

In addition to the representation problems outlined in [14], we further claim that there is in fact a *bias* that arises when applying proportionality metrics on count-based data, which is a result of added variability from the randomness of the sequencing process. In this paper, we quantify this bias and provide an alternative model-based means of estimating proportionality metrics in count-based compositional data.

## Methods

### Bias

In this section we outline the inherent bias in estimating metrics of proportionality using observed counts. Suppose that we have a matrix **Y**_*n×*(*J*+1)_ containing counts of *J* + 1 features (e.g. genes, species, etc.) from *n* samples. Let the entries of **Y** be *y*_*ij*_, representing the observed count of feature *j* in sample *i*. We seek to compare the result of plugging counts *y*_*ij*_ (or equivalently observed proportions 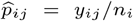 into *v, ϕ*, and *ρ* against the true (unknown) values which instead take *unobserved* proportions *p*_*ij*_ as arguments to these metrics. In each case, there is no exact closed form for the bias, and we instead use an approximation based on a second-order Taylor series expansion. We summarize the results here, but full derivations can be found in Section S2. In each case, it is clear that the bias is more pronounced when the read depths are small, on average. However, it is also evident that the amount of variation in the read depths itself plays into the bias, with more variability leading to a larger discrepancy between the empirical and true values of the metrics. The underlying distribution of the unobserved proportion vector **p**_*i*_ = (*p*_*i*1_, …, *p*_*i*(*J*+1)_) is also relevant.

First, we consider *v*(*y*_*ij*_, *y*_*ik*_), which can be approximated as:

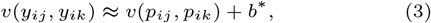

where,

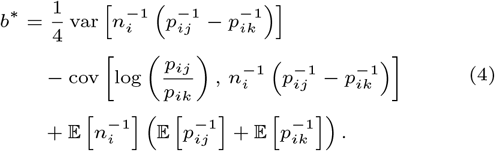

As expected, the expectation of 1*/n*_*i*_ is an important factor, however, the expectations of 1*/p*_*ij*_ and 1*/p*_*ik*_ are also present, suggesting that the discrepancy will be more pronounced in less abundant features. The bias for *ϕ* can be expressed as:

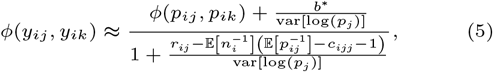

where

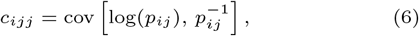

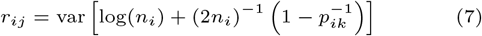

Finally, we consider the analogous result for *ρ*:

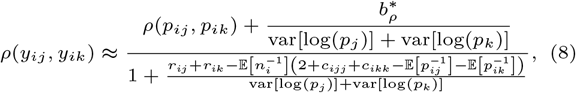

where,

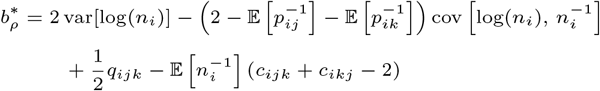

and

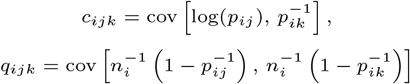

We can once again see that the bias is less pronounced when the mean of the read depths *n*_*i*_ is large. However, the variation of *n*_*i*_ as well as distribution of the proportions **p**_*i*_ are also important. This suggests that there could still be bias present for estimates for certain feature pairs even when the read depths are large.

To numerically investigate these biases, we run a small simulation from a multinomial logit-normal distribution (see the next section for the definition of this distribution). We simulate *J* + 1 = 7 features on *n* = 2000 samples, with 1000 simulation replications. In each replication, the estimated proportionality metrics on the raw counts *v*(*y*_*ij*_, *y*_*ik*_), *ϕ*(*y*_*ij*_, *y*_*ik*_), and *ρ*(*y*_*ij*_, *y*_*ik*_) are calculated (after imputing zeros using Bayesian-multiplicative replacement [22], if necessary) and compared to their respective true values *v*(*p*_*ij*_, *p*_*ik*_), *ϕ*(*p*_*ij*_, *p*_*ik*_), and *ρ*(*p*_*ij*_, *p*_*ik*_). Additionally, a bias-corrected version of the estimator is calculated by adjusting the empirical estimates using Equations 3, 5, and 8.

Figure 1 shows the difference between the estimated values and the true values for a single element of the matrix. It is evident that all of *v, ϕ*, and *ρ* exhibit strong biases when calculated on the raw counts. In all cases, the sampling distribution of the corrected estimator is approximately centered around zero. Results for the full matrix can be seen in Figures S1-S3.

**Fig. 1.**
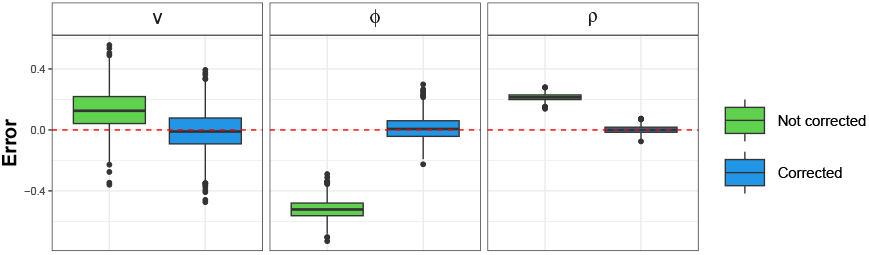
Estimation error for a single entry of *v, ϕ* and *ρ* calculated using observed counts in 2000 simulation replications. The green is not corrected for bias, the blue is corrected for bias using Equations 3, 5, and 8.

It should be noted that this bias-correction approach is not feasible in practice, as it would require knowledge of the true population parameters, which happen to be known in this simulation. As an alternative, we propose directly modelling the distribution of the proportions **p**_*i*_. The parameters of this distribution then define the values of *v, ϕ*, and *ρ*, allowing the proportionality metrics to be estimated free from the variation induced by the sequencing process itself.

### Model

We proceed in the context of the multinomial logit-normal model, posited by [30]. This model has been extremely popular in statistical modelling of the microbiome [29, 24, 17]. In this formulation, we assume that the count vector for sample *i*, denoted by **Y**_*i*_ = (*y*_*i*1_, …, *y*_*i*(*J*+1)_), is distributed as:

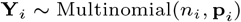

where *J* + 1 is the total number of features observed among all samples and 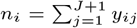 is the read depth. The proportion vector for individual *i* is assumed to follow the logit-normal distribution, which is characterized by the inverse additive log-ratio (ALR) transformation:

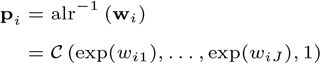

where 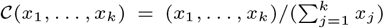 is the closure operation. Each **w**_*i*_ = (*w*_*i*1_, …, *w*_*iJ*_) is an unobserved, latent vector assumed to be multivariate-normal:

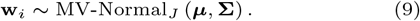

We set the (*J* + 1)^*th*^ feature as the reference feature in the ALR transformation (without loss of generality; any feature could be moved to that column). The read depths themselves are random variables on which we assume a log-normal distribution:

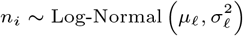

The log-normal distribution is appropriate given that the read depth distribution in a dataset can span multiple orders of magnitude. This assumption has been used in previous literature [16].

To handle the possibility of a large number of features *J* + 1 relative to the sample size *n*, we apply the Graphical Lasso (GLasso) penalty [8] to the multivariate normal log-likelihood for the **w**_*i*_ vectors:

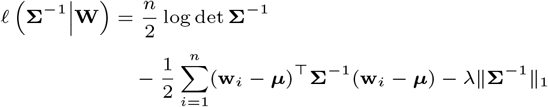

where || · ||_1_ denotes the *L*_1_-norm, and *λ* ≥ 0 is the penalty parameter. This penalization is very important in guarding against spurious associations when the number of features is large.

### Estimating proportionality metrics

In the logit-normal framework, the variation matrix elements have a convenient form, namely:

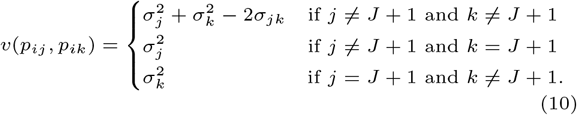

For *ϕ* and *ρ* we also need to estimate the covariance matrix of the log-proportions, which we denote by **Ω** = var[log(**p**)]. The relationship between the covariance of the log-proportions is given by **Σ** = **FΩF**^⊤^, where **F** = [**I**_*J*_, −**1**_*J*_], with **I**_*J*_ representing the *J × J* identity matrix, and **1**_*J*_ representing a *J*-dimensional vector of ones. In the logit-normal distribution there is no closed-form expression for the matrix **Ω**. The additive log-ratio variance **Σ** does not itself admit a unique **Ω** [1]; the mean of the distribution is also required to determine **Ω**.

Consequently, to estimate the log-proportion variance **Ω** from ***μ*** and **Σ**, we use a second-order Taylor approximation [10] based on the transformation from the *w*_*ij*_ scale to the log(*p*_*ij*_) scale. We can approximate the covariance of the log-proportion vector as:

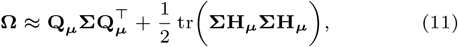

where,

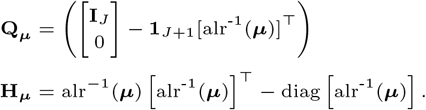

Estimates of *ϕ*, and *ρ* can then be obtained by plugging the approximations from Equations 10 and 11 into their respective definitions.

### Fitting the model

To fit the logit-normal multinomial model with graphical lasso penalty applied to the ALR-scale precision matrix, we perform maximum likelihood estimation through an implementation of the EM-algorithm posited in [11, 27]. Model selection (through the choice of tuning parameter *λ*) is performed using the Extended Bayesian Information Criterion (EBIC) [4]. The steps of the algorithm can be found in Section S4.

### Simulation study

We performed an extensive simulation study to compare the performance of model-based vs. empirical estimates. We focus on *ρ* in the simulations since it is the most interpretable metric, and we can fix it to zero for certain feature pairs allowing precision-recall analysis (whereas the null for *v* and *ϕ* is infinity). We also include an additional estimator, which uses empirical estimates of ***μ*** and **Σ**, based on simply taking the additive log-ratio of the observed counts and calculating the sample means and variances of the features. These parameter estimates are then inserted into Equation 11. We will refer to this latter estimator as the “plugin” estimator, and does not require running maximum likelihood estimation.

Our simulation parameters are based on an initial multinomial logit-normal model fitted to single-cell RNA-sequencing data from a mouse embryonic stem-cell dataset from Buettner et al. [2]. The model parameters ***μ*, Ω, Σ**, *μ*_*ℓ*_, and *σ* _*ℓ*_, are estimated from the data and are then used as the ground truth in the simulations. See Section S5 for more details about the simulation procedures.

Several scenarios are considered. The first scenario uses all the parameter values that estimated from the Buettner dataset. This scenario allows us to compare the estimated values to the true values used in the simulation using the root mean squared error (RMSE).

The second scenario modifies **Ω** to be a sparse matrix, which allows some of the true values of *ρ*(*p*_*ij*_, *p*_*ik*_) to be zero. This lets us ascertain the ability of the model-based and empirical estimates to find the non-zero elements of *ρ*(*p*_*ij*_, *p*_*ik*_). Since neither the model-based nor empirical estimators will set values exactly to zero, a range of thresholds is applied so that estimates whose *ρ* value is greater (in absolute value) than the threshold are considered to be estimated as non-zero. This allows us to calculate the area under the precision-recall curve to compare the model-based and empirical estimates.

The third scenario considers model misspecification. The motivation is to simulate values of the **w**_*i*_ vectors from a distribution that is non-Gaussian to see how estimates of *ρ* are affected by a misspecified model. To do this, we instead simulate **w**_*i*_ from a multivariate non-central *t*-distribution with mean ***μ***, covariance **Σ**, and degrees of freedom parameter equal to 2.1. This allows the distribution of the **w**_*i*_ terms to have heavier tails than the normal distribution as well as skewness. The rest of the simulation procedure is the same as in the second scenario. Precision-recall curves are used to compare performance.

### Data application

To demonstrate the applicability of the proportionality metric estimation technique introduced in this paper, we apply the methods to two datasets—one is a single-cell RNA-sequencing dataset, and the other is a metagenomic dataset.

The first dataset is a mouse embryonic stem-cell (mESC) single-cell RNA-sequencing dataset from [2]. Flow cytometry was used to sort by cell-type, with cells sorted into G1, S, and G2M cell cycle stages. The original dataset contained 96 samples and 38,390 genes. The goal of our analysis is to focus on genes that were previously shown to be differentially expressed among cell-types. In this analysis, we consider 570 genes associated with cell-cycle based on GO annotations; this list was provided in the original publication. We took a further subset of genes by filtering out genes having greater than 20% dropouts in at least one of the cell-cycle stages. There were 303 genes remaining after filtering. This analysis allows investigation of whether gene-gene associations differ between cell-cycle stages in a subset of genes that were previously known to be related to cell-cycle.

The second dataset comprises shotgun metagenomic sequencing information derived from stool samples procured from participants with pediatric-onset (symptom onset *<* 18 years of age) multiple sclerosis (MS) and unaffected controls [20, 19]. The goal of the study was to determine how suppression of dietary fibre fermentation can induce inflammation in MS. Data were collected through the Canadian Pediatric Demyelinating Disease Network; all participants were under 22 years of age at the time of stool sample procurement. There were 17 MS participants (14 female) and 20 unaffected controls (16 female) who provided a stool sample. For our analyses, we proceed at the genus level; initially there were 617 genera available in the dataset. We filtered out genera with greater than 10% zeros, leaving 296 genera. Due to the very small sample size of this dataset, we took a further subset of the 100 most abundant remaining genera. This facilitated more stable estimates of the model parameters in the small sample size case. The analysis focuses on which genus pairs have differing associations between MS participants and unaffected controls.

## Results

### Simulation results

Results from the first simulation can be seen in Section S6. The goal of this initial simulation is to investigate the estimation accuracy of *ρ* and the elements of **Ω**. In Figure S4, the RMSE is shown for empirical and model-based estimation of *ρ*. It is clear that there is a substantial improvement in model-based estimates in most cases, especially when the number of features *J* is large, which is an important guard against finding spurious correlations. We show similar results for off-diagonal elements of **Ω** in Figure S5. The RMSE patterns for **Ω** are congruent with those of *ρ*.

Next we show the results from the simulation scenario considering sparsity among the *ρ* values. Results can be seen in Figure 2, where we compare the distributions of the AUPRC for the different estimates. In all scenarios, the model-based estimates outperform the empirical estimates. The difference in performance is especially pronounced when the number of features *J* is large. Intriguingly, even the plugin estimates of *ρ* outperform the empirical estimates. This is also an attractive option, as it does not require running maximum likelihood estimation.

**Fig. 2.**
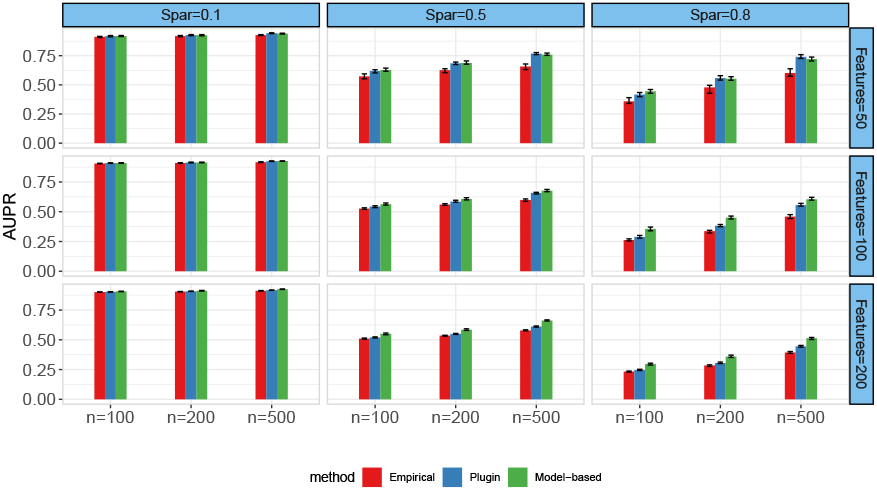
Area under the precision-recall curve of *ρ* estimation methods in simulation data. Error bars represent 25th and 75th percentiles over 50 simulation replications. Simulations vary over sparsity level (e.g. proportion of true *ρ* values equal to 0) and number of features *J*.

Finally, we present the results from repeating the AUPRC simulation, but under severe model misspecification (non-central *t*-distribution in place of Gaussian). These results can be seen in Figure S6. Though the differences between the AUPRC are more subdued in this case, there is still a clear advantage for the model-based estimator over the empirical estimator. There is, again, a slight advantage for the plugin estimator over the empirical estimator. This is an encouraging result, as it demonstrates the utility of our estimator for *ρ* even if the form of the assumed model is not correct.

All of the simulations show strong evidence of the superiority of the model-based estimates over the empirical estimates for *ρ* elements. Thus, the model-based estimates can be used to guard against spurious correlations in when considering the proportionality metric *ρ*.

### Data application results

In this section, we outline the results from the two data analyses described in the Methods section. First we discuss the results from the murine single-cell RNA-seq dataset. All gene descriptions were obtained from GeneCards [26].

We consider the estimated *ρ* values for each of the pairs of the 303 genes retained after the aforementioned filtering steps. Estimates were obtained using both the empirical estimator and the model-based estimator. We compare the top 100 gene-gene associations (in absolute value) detected using the empirical and model-based estimators. In the G1 phase there were 23 gene pairs in common, in G2M phase there were 24 gene pairs in common, and in S phase there was only 1 gene pair in common (Figure S7). Relatedly, the empirical and model-based estimates are compared in Figure S8. This shows the attenuation towards the null for the model-based estimates, which is an important guard against spurious correlations.

These comparisons between the empirical and model-based estimates show how different the results can be using the empirical estimator for *ρ* and highlights the importance of correcting the inherent biases in proportionality metrics applied to count data.

The top 100 gene pairs within each cell stage are shown in Tables S4-6. Additionally, Table 1 summarizes the top differences (in absolute value) in *ρ* values between phases G1 and G2M phases; more extensive tables showing the top 100 gene pairs for each of the cell phase comparisons are shown in Tables S1-S3.

**Table 1.**
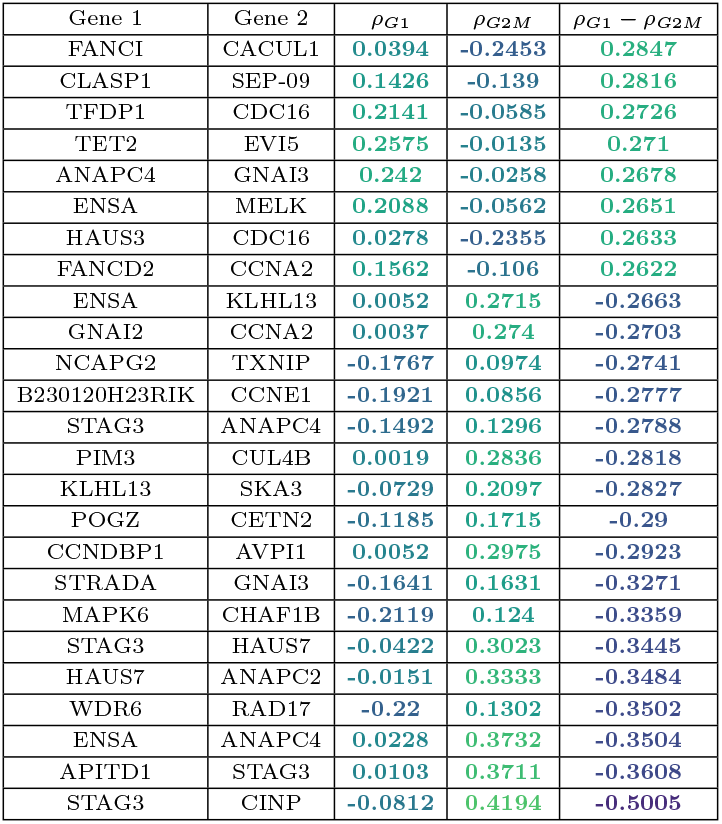
Top 25 gene pairs whose ρ value differs between G1 and G2M stages.

In the G1 vs. G2M comparison, the largest *ρ* difference is seen between genes STAG3 and CINP, with *ρ*_*G*1_ − *ρ*_*G*2*M*_ = −0.5005. STAG3 is a encodes a protein involved in the regulation of the cohesion of sister chromatids during cell division; CINP is part of the DNA replication complex and binds chromatin in the G1 phase.

In the G1 vs. S comparison (Table S2), the four largest differences all involve the ARHGEF2 gene, which is implicated in Rho-GTPase activation. The genes its *ρ* values differ with the most between G1 and S are ATM, STAG2, CHEK1, and ANAPC4, all of whose expression is much more positively correlated with ARHGEF2 in S phase compared to G1 phase.

In the G2M vs. S comparison (Table S3), ARHGEF2 is again involved in several of the top *ρ* differences; namely, with ATM, CHEK1, STAG2, and ANAPC4, again with much stronger positive correlation in S phase compared to G1 phase. Another notable difference is for ANAPC4 and PPM1G, which are slightly negatively correlated in G2M phase and positively correlated in S phase. ANAPC4 is involved in promotion of the metaphase-anaphase transition, and PPM1G is related to negative regulation of cell stress response.

Next we compare *ρ* values between MS participants and unaffected controls at the genus level. Table S7 shows the highest *ρ* values (in absolute value) in both MS and unaffected controls. The top 25 differences between MS and controls are shown in Table 2; a larger table with more genus pairs is shown in Table S8. Genera appearing among the top 25 differences that were previously shown to be associated with MS include Megasphaera [12]; Acidaminococcus [13]; Ruminococcus [3]; Dialister, Lachnospira, and Adlercreutzia [28]; Bacteroides and Prevotella [21, 28]; Lactobacillus [5], Adlercreutzia [28, 5]; and Eubacterium [23].

**Table 2.** Top 25 genus pairs whose ρ value differs between MS (ρ_MS_) and control samples (ρ_C_).

| Genus 1 | Genus 2 | $\rho_{MS}$ | $\rho_C$ | $\rho_{MS} - \rho_C$ |
| --- | --- | --- | --- | --- |
| UBA1417 | BacteroidesB | -0.2295 | 0.2525 | -0.4819 |
| EubacteriumR | Prevotella | -0.203 | 0.2838 | -0.4868 |
| Dialister | UBA1417 | -0.071 | 0.4202 | -0.4913 |
| UBA11774 | Acidaminococcus | -0.0573 | 0.4377 | -0.495 |
| RuminiclostridiumE | LactobacillusB | -0.1886 | 0.3084 | -0.4971 |
| Senegalimassilia | Adlercreutzia | -0.2275 | 0.2735 | -0.501 |
| RuminiclostridiumC | CAG-83 | 0.0213 | 0.5311 | -0.5098 |
| PeH17 | BacteroidesB | -0.2355 | 0.278 | -0.5135 |
| Adlercreutzia | Dialister | -0.293 | 0.2326 | -0.5256 |
| RuminiclostridiumE | Acidaminococcus | -0.0328 | 0.4996 | -0.5323 |
| Acidaminococcus | CAG-177 | -0.0945 | 0.4461 | -0.5406 |
| Megasphaera | RuminococcusC | -0.0671 | 0.4807 | -0.5478 |
| Megasphaera | CAG-127 | -0.0442 | 0.5057 | -0.5499 |
| Acidaminococcus | EubacteriumR | -0.2135 | 0.3547 | -0.5683 |
| Megasphaera | CAG-180 | -0.1967 | 0.3853 | -0.5819 |
| Acidaminococcus | Adlercreutzia | -0.1906 | 0.413 | -0.6036 |
| EubacteriumR | Dialister | -0.2587 | 0.3482 | -0.6069 |
| Megasphaera | CAG-177 | -0.1881 | 0.4253 | -0.6135 |
| Megasphaera | RuminococcusD | -0.1457 | 0.4686 | -0.6143 |
| Acidaminococcus | CAG-180 | -0.2094 | 0.4055 | -0.6149 |
| RuminiclostridiumE | Megasphaera | -0.0811 | 0.5342 | -0.6154 |
| UBA11774 | Lachnospira | -0.1046 | 0.5132 | -0.6178 |
| Acidaminococcus | RuminococcusC | -0.0643 | 0.563 | -0.6273 |
| Acidaminococcus | CAG-127 | -0.0385 | 0.6221 | -0.6606 |
| UBA11774 | Dialister | -0.1607 | 0.5012 | -0.6619 |

In all of the top 25 genus pairs, the estimated *ρ*-values are greater in controls compared to MS. The values for controls are positive and the MS values are either close to zero or negative. This trend is present in the top 100 genus pairs as well, with a few exceptions. In Figure S9, we see that there are only 25 genus pairs common in the top 100 pairs for MS and unaffected controls. This analysis has uncovered a strong disruption in community-level dynamics in the MS participants among a number of aforementioned genera whose abundances were already known to be related to MS.

## Conclusion

We have demonstrated that empirical estimates of metrics of proportionality in count-based platforms can lead to bias. Though the bias is mitigated when the mean read depths is large, there may still exist a bias in less-abundant features regardless. We therefore designed a multinomial logit-normal model to calculate model-based estimates of the proportionality metrics. We showed that, in an extensive simulation study, the model-based estimates outperformed the empirical estimates, even in the case of model misspecification. Additionally, a simple plugin estimator outperformed empirical estimates, which could be useful in the case where a user does not have the computational resources to obtain model estimates through the maximum likelihood estimator. Importantly, empirical vs. model-based estimates differed in a way that could drastically change results regarding which features are most strongly correlated with one another.

One limitation of our approach is that the model does not differentiate between structural and sampling zeros, hence the need for filtering out features with many zero counts. Though this could be remedied in the model by introducing zero-inflated parameters as done in [31, 16], this introduces a new problem regarding the interpretation of the proportionality metrics, which are not defined for zero-valued arguments. One could conceivably define these metrics conditional on non-zero values, but this would ignore valid information contained in the probability of a zero proportion. To address this, zero-inflated versions of Kendall’s *τ* and Spearman’s *ρ* could be used in the context of a zero-inflated model. This is left for future work.

## Competing interests

H.T. has, in the last five years, received research support from the Canada Research Chair Program, the National Multiple Sclerosis Society, the Canadian Institutes of Health Research, the Multiple Sclerosis Society of Canada, the Multiple Sclerosis Scientific Research Foundation and the EDMUS Foundation (‘Fondation EDMUS contre la sclérose en plaques’). In addition, in the last five years, has had travel expenses or registration fees prepaid or reimbursed to present at CME conferences from the Consortium of MS Centres (2018, 2023), the Canadian Neurological Sciences Federation (2023), National MS Society (2018, 2022), ECTRIMS/ ACTRIMS (2017-2023), American Academy of Neurology (2019). Speaker honoraria are either declined or donated to an MS charity or to an unrestricted grant for use by H.T.’s research group.

## Acknowledgments

K.M. acknowledges support from the Natural Sciences and Engineering Research Council of Canada (NSERC Discovery Grant: RGPIN-2021-03634) as well as an NSERC Undergraduate Student Research Award for N.O. The MS study was supported by funding from Canada Research Chair (PI: H.A.), the Multiple Sclerosis Society (PI: H.A.), and the Multiple Sclerosis Scientific and Research Foundation (#EGID: 2636; PI: H.T.). These latter funding sources were not involved in the study design, the collection, analysis, and interpretation of the data, or in the decision to submit this article for publication. We are grateful for all the participants’ involvement, especially children and teenagers with MS and their parents. We are also grateful to all the investigators and study teams at each site involved in the Canadian Paediatric Demyelinating Disease Network study, without whom this study would not have been possible. We acknowledge the important contributions of the H.A. team (University of Manitoba) and H.T. team (University of British Columbia); Thomas Duggan in facilitating study set-up, coordination and data collection; Bonnie Leung for study coordination; Michael Sargent (Department of Internal Medicine, and the University of Manitoba IBD Clinical and Research Centre laboratory) for managing the stool biobank, and Jessica D. Forbes (University of Toronto) for assisting with the original grant.

## Data availability statement

The mouse embryonic stem cell dataset is publicly available under accession number E-MTAB-2512. The pediatric-onset MS dataset cannot be shared publicly for privacy of participants. The authors can be contacted for data access; requests will be assessed on a case-by-case basis, based on the scientific rigor of the research question.

## Notes

### Competing Interest Statement

Helen Tremlett has, in the last five years, received research support from the Canada Research Chair Program, the National Multiple Sclerosis Society, the Canadian Institutes of Health Research, the Multiple Sclerosis Society of Canada, the Multiple Sclerosis Scientific Research Foundation and the EDMUS Foundation (Fondation EDMUS contre la sclerose en plaques). In addition, in the last five years, has had travel expenses or registration fees prepaid or reimbursed to present at CME conferences from the Consortium of MS Centres (2018, 2023), the Canadian Neurological Sciences Federation (2023), National MS Society (2018, 2022), ECTRIMS/ ACTRIMS (2017-2023), American Academy of Neurology (2019). Speaker honoraria are either declined or donated to an MS charity or to an unrestricted grant for use by the HT research group.

