## Supplementary material for "Proportionality-based association metrics in count compositional data"

#### 1 Introduction

We model an $N \times (J+1)$ matrix $\mathbf{Y}$ containing counts corresponding to some kind of biological sequencing platform. The entry in row $i$ and column $j$ is denoted by $y_{ij}$. These data are assumed *compositional*, meaning that the magnitude of each $y_{ij}$ is not itself interpretable; it can only be interpreted relative to other counts in the same row of the matrix, by means of ratios or log-ratios for example. Each row of $\mathbf{Y}$ is denoted by $\mathbf{Y}_i$ and represents the sequencing counts of $J+1$ categories (e.g. taxa, RNA reads) for each individual $i = 1, \ldots, N$. To represent the sequencing process, row $i$ is modelled using a multinomial distribution, so that:

$$\mathbf{Y}_i | n_i, \mathbf{p}_i \sim \text{Multinomial}\left(n_i, \mathbf{p}_i\right)$$

where $\mathbf{p}_i = (p_{i1}, \ldots, p_{i(J+1)})^\top$ is a vector specifying the unknown proportions of the $J+1$ categories for individual $i$. $n_i = \sum_{j=1}^{J+1} y_{ij}$ is the sequencing depth (read depth) of sample $i$. Firstly, we assume that $\mathbf{p}_i$ has an associated probability distribution:

$$\mathbf{p}_i \sim F_{\mathbf{p},\boldsymbol{\theta}}$$

for some distribution function $F_{\mathbf{p},\boldsymbol{\theta}}$ defined by a vector of parameters $\boldsymbol{\theta}$. Note that the choice of family for $F_{\mathbf{p},\boldsymbol{\theta}}$ is restricted to those families whose support lies in the unit simplex—in other words, $\sum_{j=1}^{J+1} p_{ij} = 1$. Examples include the Dirichlet or the logit-normal distributions.

Additionally, we assume $n_i$, the number of sequencing reads for sample $i$, to be random. Though $n_i$ is a positive count, there is precedent in recent literature to approximately model $n_i$ using a log-normal distribution. For now, we'll assume a generic probability distribution for $n_i$:

$$n_i \sim F_{n,\boldsymbol{\gamma}}$$

where $\boldsymbol{\gamma}$ is a parameter vector that defines the distribution for $n_i$ under this family. Furthermore, assume that $n_i \perp\!\!\!\perp \mathbf{p}_i$. This assumption is fair given that the read depth $n_i$ is determined by technical artifacts in the sequencing process and should not be dependent on the underlying proportions $\mathbf{p}_i$.

#### 2 Variation matrix estimation bias

In compositional data, we are often interested in calculating the *variation matrix*, whose elements can be interpreted as measures of proportionality. We denote the $J \times J$ variation matrix for individual $i$ as $\mathbf{V}^{(i)}$. The variation matrix element in row $j$ and column $k$ is defined as:

$$v(y_{ij}, y_{ik}) = \text{var}\left[\log\left(\frac{y_{ij}}{y_{ik}}\right)\right] \tag{1}$$

If $y_{ij} \propto y_{ik}$, then we have that $v(y_{ij}, y_{ik}) = 0$. Thus $v(y_{ij}, y_{ik})$ can be used as a measure of association by considering its proximity to zero. This measure therefore falls under the class of *proportionality-based association metrics*. Furthermore, because of the scale invariance property of log-ratios, this measure of association is appropriate for use on compositional data.

There is one big caveat in calculating the empirical variation matrix elements on count-based compositional data. Though the variation matrix was posited as a means of measuring association in compositional data, it was meant for *continuous* measures. Lovell et al. (2020) showed that metrics of proportionality cannot be exactly represented in count-based data. In this section, we build on this argument, and show that the using the empirical variation matrix elements to estimate the true log-ratio variance applied to the unobserved proportions $\mathbf{p}_i$ is, in fact, biased.

The quantity we wish to estimate is the variation elements on the true (unobserved) proportions:

$$v(p_{ij}, p_{ik}) = \text{var}\left[\log\left(\frac{p_{ij}}{p_{ik}}\right)\right], \tag{2}$$

Since these are unobserved quantities, one could simply calculate (2) on the estimated proportions $\widehat{\mathbf{p}}_i$, which is equivalent to calculating (1) on the observed counts $y_{ij}$. The bias is due to the fact that the log-ratio variance on the observed counts is, in essence, the *marginal* variance, which includes the inherent variation in $n_i$ and $p_{ij}$. To illustrate the bias in this procedure, we begin by decomposition the marginal log-ratio variance of the counts as:

$$\operatorname{var}\left[\log\left(\frac{y_{ij}}{y_{ik}}\right)\right] = \operatorname{var}\left[\mathbb{E}\left(\log\left(\frac{y_{ij}}{y_{ik}}\right) \,\middle|\, n_i, \mathbf{p}_i\right)\right] + \mathbb{E}\left[\operatorname{var}\left(\log\left(\frac{y_{ij}}{y_{ik}}\right) \,\middle|\, n_i, \mathbf{p}_i\right)\right] \tag{3}$$

Thus, we need the expectation and variance of the log-ratio counts conditional on $n_i$ and $\mathbf{p}_i$. Firstly, for the conditional log-ratio variance, we have:

$$\begin{aligned}
\operatorname{var}\left[\log\left(\frac{y_{ij}}{y_{ik}}\right) \,\middle|\, n_i, \mathbf{p}_i\right] &= \operatorname{var}\left[\frac{y_{ij}/n_i}{y_{ik}/n_i} \,\middle|\, n_i, \mathbf{p}_i\right] \\
&= \operatorname{var}\Big[\log(y_{ij}/n_i) - \log(y_{ik}/n_i) \,\Big|\, n_i, \mathbf{p}_i\Big] \\
&= \operatorname{var}\Big[\log(\widehat{p}_{ij}) - \log(\widehat{p}_{ik}) \,\Big|\, n_i, \mathbf{p}_i\Big] \\
&= \operatorname{var}\Big[\log(\widehat{p}_{ij}) \,\Big|\, n_i, \mathbf{p}_i\Big] + \operatorname{var}\Big[\log(\widehat{p}_{ik}) \,\Big|\, n_i, \mathbf{p}_i\Big] - 2\operatorname{cov}\Big[\log(\widehat{p}_{ij}), \log(\widehat{p}_{ik}) \,\Big|\, n_i, \mathbf{p}_i\Big]
\end{aligned} \tag{4}$$

where $\widehat{p}_{ij} = y_{ij}/n_i$ is the observed proportion of feature $j$ in sample $i$. For convenience of notation, let $\widehat{\mathbf{p}}_i = (\widehat{p}_{i1}, \ldots, \widehat{p}_{i(J+1)})$. In order to calculate necessary components in Equation 4, we need to obtain the conditional variance-covariance matrix of $\log(\widehat{\mathbf{p}}_i) = [\log(\widehat{p}_{i1}), \ldots \log(\widehat{p}_{i(J+1)})]$.

First, we use the fact that, under the multinomial distribution

$$\sqrt{n_i}(\widehat{\mathbf{p}}_i - \mathbf{p}_i) \xrightarrow{d} \text{MVN}\left(\mathbf{0}, \operatorname{diag}(\mathbf{p}_i) - \mathbf{p}_i\mathbf{p}_i^\top\right)$$

(Mukhopadhyay, 2016). Thus, we can approximate the variance of $\log(\widehat{\mathbf{p}}_i)$ using the delta method. To do this, we need the Jacobian of the transformation $g(\widehat{\mathbf{p}}_i) = \log(\widehat{\mathbf{p}}_i)$, which is calculated as:

$$\mathbf{J}_i = \begin{bmatrix} \frac{1}{p_{i1}} & 0 & \cdots & 0 \\ 0 & \frac{1}{p_{i2}} & \cdots & 0 \\ \vdots & \vdots & \ddots & \vdots \\ 0 & 0 & 0 & \frac{1}{p_{i(J+1)}} \end{bmatrix}$$

The conditional variance-covariance matrix of the log-transformed can then be approximated by the delta method as:

$$\begin{aligned}
\operatorname{var}\left[\log(\widehat{\mathbf{p}}_i) \mid n_i, \mathbf{p}_i\right] &\approx \frac{1}{n_i}\mathbf{J}_i\Big(\operatorname{diag}(\mathbf{p}_i) - \mathbf{p}_i\mathbf{p}_i^\top\Big)\mathbf{J}_i^\top \\
&= \frac{1}{n_i}\Big(\mathbf{J}_i - \mathbf{1}_{J+1}\mathbf{1}_{J+1}^\top\Big),
\end{aligned} \tag{5}$$

where $\mathbf{1}_{J+1}$ is a $(J+1) \times 1$ vector of ones. Then, by plugging the necessary elements of the result of Equation 5 back into Equation 4, we obtain the following expression for the element in row $i$ and column $j$ of the variation matrix, conditional on $n_i$ and $\mathbf{p}_i$:

$$\text{var}\left[\log\left(\frac{y_{ij}}{y_{ik}}\right) \,\middle|\, n_i, \mathbf{p}_i\right] = \frac{1}{n_i}\left[\frac{1}{p_{ij}} + \frac{1}{p_{ik}}\right] \tag{6}$$

Next, we need to marginalize out both $n_i$ and $\mathbf{p}_i$. To do so, we employ a second-order Taylor series approximation to the conditional expectation of $\log(y_{ij})$:

$$\mathbb{E}\left[\log(y_{ij}) \mid n_i, \mathbf{p}_i\right] \approx \log(n_i) + \log(p_{ij}) + \frac{1}{2n_i}\left(1 - \frac{1}{p_{ij}}\right). \tag{7}$$

Thus, the conditional expectation of the log-ratio is given by:

$$\mathbb{E}\left[\log\left(\frac{y_{ij}}{y_{ik}}\right) \,\middle|\, n_i, \mathbf{p}_i\right] = \mathbb{E}\left[\log(y_{ij}) \mid n_i, \mathbf{p}_i\right] - \mathbb{E}\left[\log(y_{ik}) \mid n_i, \mathbf{p}_i\right] \tag{8}$$

$$\approx \log\left(\frac{p_{ij}}{p_{ik}}\right) - \frac{1}{2n_i}\left(\frac{1}{p_{ij}} - \frac{1}{p_{ik}}\right) \tag{9}$$

Now, to proceed with the marginalization, we consider both components in the right-hand side of Equation 3. The first term is:

$$\begin{aligned}
\text{var}\left[\mathbb{E}\left(\log\left(\frac{y_{ij}}{y_{ik}}\right) \,\middle|\, n_i, \mathbf{p}_i\right)\right] &\approx \text{var}\left[\log\left(\frac{p_{ij}}{p_{ik}}\right) - \frac{1}{2n_i}\left(\frac{1}{p_{ij}} - \frac{1}{p_{ik}}\right)\right] \\
&= \text{var}\left[\log\left(\frac{p_{ij}}{p_{ik}}\right)\right] + \text{var}\left[\frac{1}{2n_i}\left(\frac{1}{p_{ij}} - \frac{1}{p_{ik}}\right)\right] \\
&\quad - 2\,\text{cov}\left[\log\left(\frac{p_{ij}}{p_{ik}}\right), \frac{1}{2n_i}\left(\frac{1}{p_{ij}} - \frac{1}{p_{ik}}\right)\right] \\
&= v(p_{ij}, p_{ik}) + \text{var}\left[\frac{1}{2n_i}\left(\frac{1}{p_{ij}} - \frac{1}{p_{ik}}\right)\right] - 2\,\text{cov}\left[\log\left(\frac{p_{ij}}{p_{ik}}\right), \frac{1}{2n_i}\left(\frac{1}{p_{ij}} - \frac{1}{p_{ik}}\right)\right]
\end{aligned} \tag{10}$$

Likewise, the second term is:

$$\begin{aligned}
\mathbb{E}\left[\text{var}\left(\log\left(\frac{y_{ij}}{y_{ik}}\right) \,\middle|\, n_i, \mathbf{p}_i\right)\right] &= \mathbb{E}\left[\frac{1}{n_i}\left(\frac{1}{p_{ij}} + \frac{1}{p_{ik}}\right)\right] \\
&= \mathbb{E}\left[\frac{1}{n_i}\right]\left(\mathbb{E}\left[\frac{1}{p_{ij}}\right] + \mathbb{E}\left[\frac{1}{p_{ik}}\right]\right)
\end{aligned} \tag{11}$$

Plugging 10 and 11 back into 3 characterizes the bias:

$$\begin{aligned}
\operatorname{var}\left[\log\left(\frac{y_{ij}}{y_{ik}}\right)\right] &= \operatorname{var}\left[\mathbb{E}\left(\log\left(\frac{y_{ij}}{y_{ik}}\right) \,\middle|\, n_i, \mathbf{p}_i\right)\right] + \mathbb{E}\left[\operatorname{var}\left(\log\left(\frac{y_{ij}}{y_{ik}}\right) \,\middle|\, n_i, \mathbf{p}_i\right)\right] \\
&\approx v(p_{ij}, p_{ik}) + \operatorname{var}\left[\frac{1}{2n_i}\left(\frac{1}{p_{ij}} - \frac{1}{p_{ik}}\right)\right] - 2\operatorname{cov}\left[\log\left(\frac{p_{ij}}{p_{ik}}\right), \frac{1}{2n_i}\left(\frac{1}{p_{ij}} - \frac{1}{p_{ik}}\right)\right] \\
&\quad + \mathbb{E}\left[\frac{1}{n_i}\right]\left(\mathbb{E}\left[\frac{1}{p_{ij}}\right] + \mathbb{E}\left[\frac{1}{p_{ik}}\right]\right) \\
&= v(p_{ij}, p_{ik}) + b^*,
\end{aligned} \tag{12}$$

where $b^*$ is the approximate bias term, and is equal to:

$$b^* = \operatorname{var}\left[\frac{1}{2n_i}\left(\frac{1}{p_{ij}} - \frac{1}{p_{ik}}\right)\right] - 2\operatorname{cov}\left[\log\left(\frac{p_{ij}}{p_{ik}}\right), \frac{1}{2n_i}\left(\frac{1}{p_{ij}} - \frac{1}{p_{ik}}\right)\right] + \mathbb{E}\left[\frac{1}{n_i}\right]\left(\mathbb{E}\left[\frac{1}{p_{ij}}\right] + \mathbb{E}\left[\frac{1}{p_{ik}}\right]\right). \tag{13}$$

This result shows that this bias term shrinks with increasing mean $n_i$, e.g. the mean read depth. However, this also demonstrates that the amount of variation in the read depth contributes to the estimation bias. Similarly, due to the presence of the expectations of $p_{ij}^{-1}$ and $p_{ik}^{-1}$, we expect that low-abundance features will be more affected by the bias, potentially even in the case where the expected read depth is reasonably large.

### 3 Bias for $\phi$ and $\rho$

The bias for $\phi$ and $\rho$ is more complicated. The true value of $\phi$ is given by:

$$\phi(p_{ij}, p_{ik}) = \frac{v(p_{ij}, p_{ik})}{\operatorname{var}[\log(p_{ij})]},$$

and likewise for $\rho$:

$$\rho(p_{ij}, p_{ik}) = \frac{2\operatorname{cov}\left[\log(p_{ij}), \log(p_{ik})\right]}{\operatorname{var}[\log(p_{ij})] + \operatorname{var}[\log(p_{ik})]}.$$

In each case, we have $\operatorname{var}[\log(p_{ij})]$ in the denominator. Here we consider the effect of using the counts to estimate this quantity, i.e. $\operatorname{var}[\log(y_{ij})]$, which can be decomposed as:

$$\operatorname{var}\left[\log\left(y_{ij}\right)\right] = \operatorname{var}\left[\mathbb{E}\left(\log\left(y_{ij}\right) \mid n_i, \mathbf{p}_i\right)\right] + \mathbb{E}\left[\operatorname{var}\left(\log\left(y_{ij}\right) \mid n_i, \mathbf{p}_i\right)\right] \tag{14}$$

The first term of the decomposition is approximated using Equation 7 as:

$$\begin{aligned}\operatorname{var}\left[\mathbb{E}\left(\log\left(y_{ij}\right) \mid n_i, \mathbf{p}_i\right)\right] &\approx \operatorname{var}\left[\log(n_i) + \log(p_{ij}) + \frac{1}{2n_i}\left(1 - \frac{1}{p_{ij}}\right)\right] \\ &= \operatorname{var}\left[\log\left(p_{ij}\right)\right] + \operatorname{var}\left[\log(n_i) + \frac{1}{2n_i}\left(1 - \frac{1}{p_{ij}}\right)\right] \\ &\quad + 2\operatorname{cov}\left[\log(p_{ij}),\ \log(n_i) + \frac{1}{2n_i}\left(1 - \frac{1}{p_{ij}}\right)\right]\end{aligned} \tag{15}$$

Likewise, the second term is given by:

$$\begin{aligned}\mathbb{E}\left[\operatorname{var}\left(\log\left(y_{ij}\right) \mid n_i, \mathbf{p}_i\right)\right] &= \mathbb{E}\left[\operatorname{var}\left(\log\left(y_{ij}\right) - \log(n_i) \mid n_i, \mathbf{p}_i\right)\right] \\ &\approx \mathbb{E}\left[\frac{1}{n_i}\left(\frac{1}{p_{ij}} - 1\right)\right] \\ &= \mathbb{E}\left[\frac{1}{n_i}\right]\left(\mathbb{E}\left[\frac{1}{p_{ij}}\right] - 1\right)\end{aligned}$$

which uses the variance calculation in Equation 5. We can then formulate the following approximation:

$$\begin{aligned}\operatorname{var}\left[\log\left(y_{ij}\right)\right] &\approx \operatorname{var}\left[\log\left(p_{ij}\right)\right] + \operatorname{var}\left[\log(n_i) + \frac{1}{2n_i}\left(1 - \frac{1}{p_{ij}}\right)\right] \\ &\quad + 2\operatorname{cov}\left[\log(p_{ij}),\ \log(n_i) + \frac{1}{2n_i}\left(1 - \frac{1}{p_{ij}}\right)\right] + \mathbb{E}\left[\frac{1}{n_i}\right]\left(\mathbb{E}\left[\frac{1}{p_{ij}}\right] - 1\right)\end{aligned} \tag{16}$$

#### 3.1 Bias for $\phi$

Using the results derived in Equations 12 and 16, as well as some algebraic manipulation, we can approximate the relationship between $\phi(y_{ij}, y_{ik})$ and $\phi(p_{ij}, p_{ik})$ as:

$$\phi(y_{ij}, y_{ik}) \approx \frac{\phi(p_{ij}, p_{ik}) + \frac{b^*}{\operatorname{var}[\log(p_j)]}}{1 + \frac{r_{ij} - \mathbb{E}\left[n_i^{-1}\right]\left(1 + c_{ijj} - \mathbb{E}\left[p_{ij}^{-1}\right]\right)}{\operatorname{var}[\log(p_j)]}}, \tag{17}$$

where $c_{ijj} = \operatorname{cov}\left[\log(p_{ij}),\ \frac{1}{p_{ij}}\right]$ and $r_{ij} = \operatorname{var}\left[\log(n_i) + \frac{1}{2n_i}\left(1 - \frac{1}{p_{ik}}\right)\right]$.

#### 3.2 Bias for $\rho$

Similarly, using the results derived in Equation 16, as well as some algebraic manipulation, we can approximate the relationship between $\rho(y_{ij}, y_{ik})$ and $\rho(p_{ij}, p_{ik})$ as:

$$\rho(y_{ij}, y_{ik}) \approx \frac{\rho(p_{ij}, p_{ik}) + \frac{2\operatorname{var}[\log(n_i)] - \left(2 - \mathbb{E}\left[p_{ij}^{-1}\right] - \mathbb{E}\left[p_{ik}^{-1}\right]\right)\operatorname{cov}\left[\log(n_i),\, n_i^{-1}\right] + \frac{1}{2}q_{ijk} - \mathbb{E}\left[n_i^{-1}\right](c_{ijk} + c_{ikj} - 2)}{\operatorname{var}[\log(p_j)] + \operatorname{var}[\log(p_k)]}}{1 + \frac{r_{ij} + r_{ik} - \mathbb{E}\left[n_i^{-1}\right]\left(2 + c_{ijj} + c_{ikk} - \mathbb{E}\left[p_{ij}^{-1}\right] - \mathbb{E}\left[p_{ik}^{-1}\right]\right)}{\operatorname{var}[\log(p_j)] + \operatorname{var}[\log(p_k)]}}, \tag{18}$$

where $c_{ijk} = \text{cov}\left[\log(p_{ij}),\, \frac{1}{p_{ik}}\right]$ and $q_{ijk} = \text{cov}\left[\frac{1}{n_i}\left(1 - \frac{1}{p_{ij}}\right),\, \frac{1}{n_i}\left(1 - \frac{1}{p_{ik}}\right)\right]$.

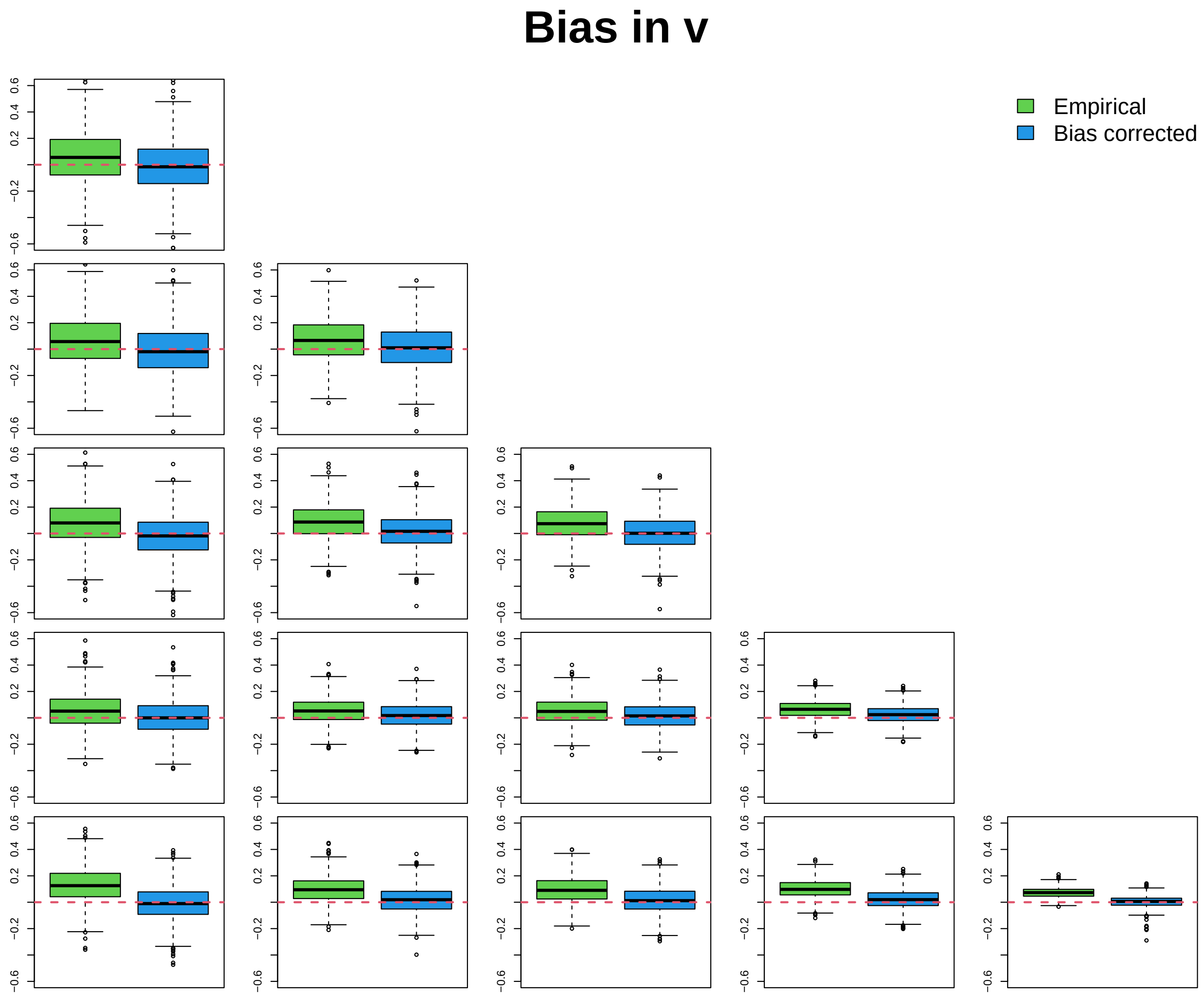


Figure S1: Bias in $v(y_{ij}, y_{ik})$ over 1000 simulation replications. The vertical axis is the estimated value minus the true value, $v(p_{ij}, p_{ik})$. Each plot corresponds to a pair $(j, k)$ with $j \neq k$. The green boxplots show the empirical estimates, and the blue boxplots are the empirical estimates adjusted by subtracting off the bias term in Equation 13.

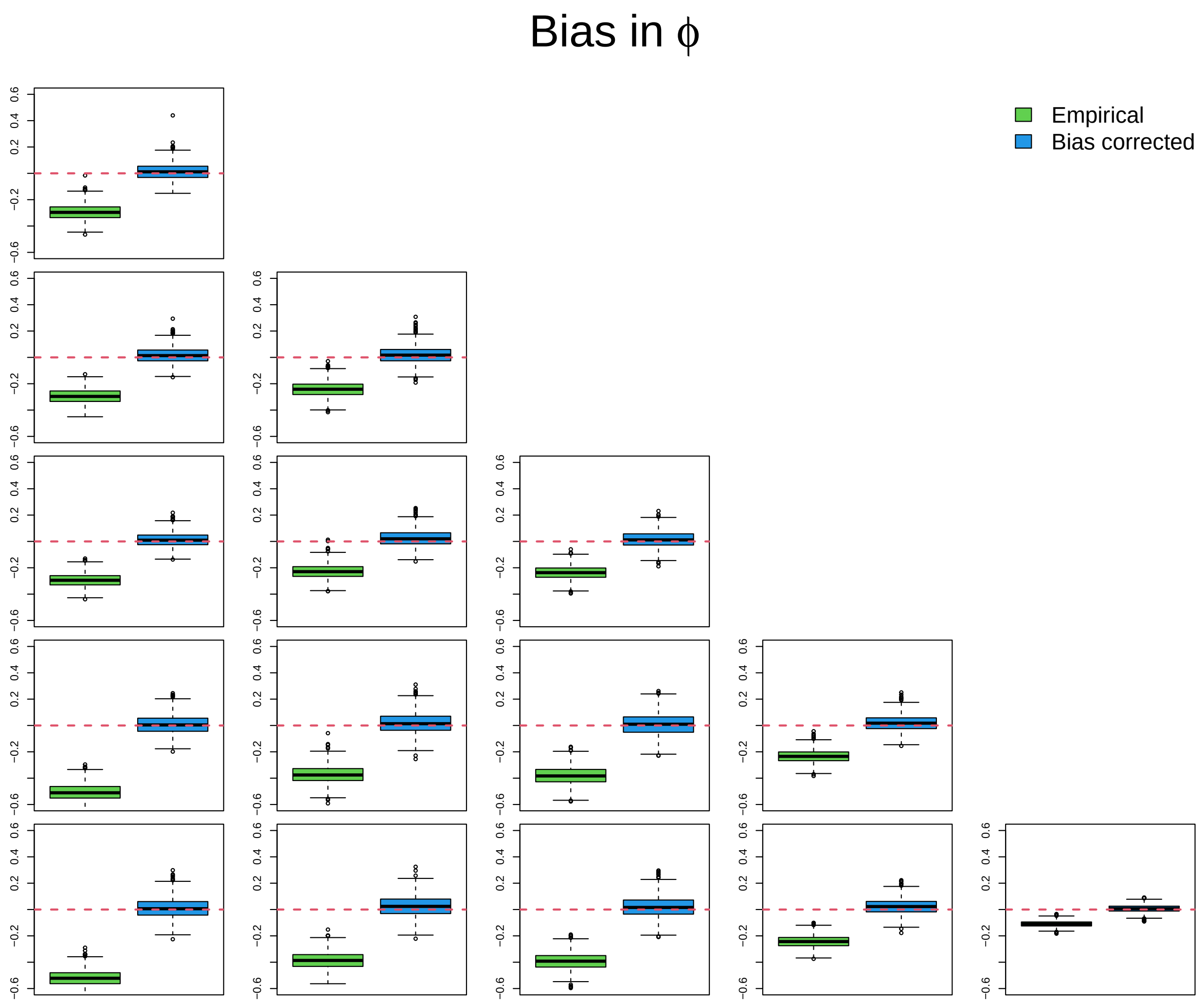


Figure S2: Bias in $\phi(y_{ij}, y_{ik})$ over 1000 simulation replications. The vertical axis is the estimated value minus the true value, $\phi(p_{ij}, p_{ik})$. Each plot corresponds to a pair $(j, k)$ with $j \neq k$. The green boxplots show the empirical estimates, and the blue boxplots are the empirical estimates adjusted using Equation 17.

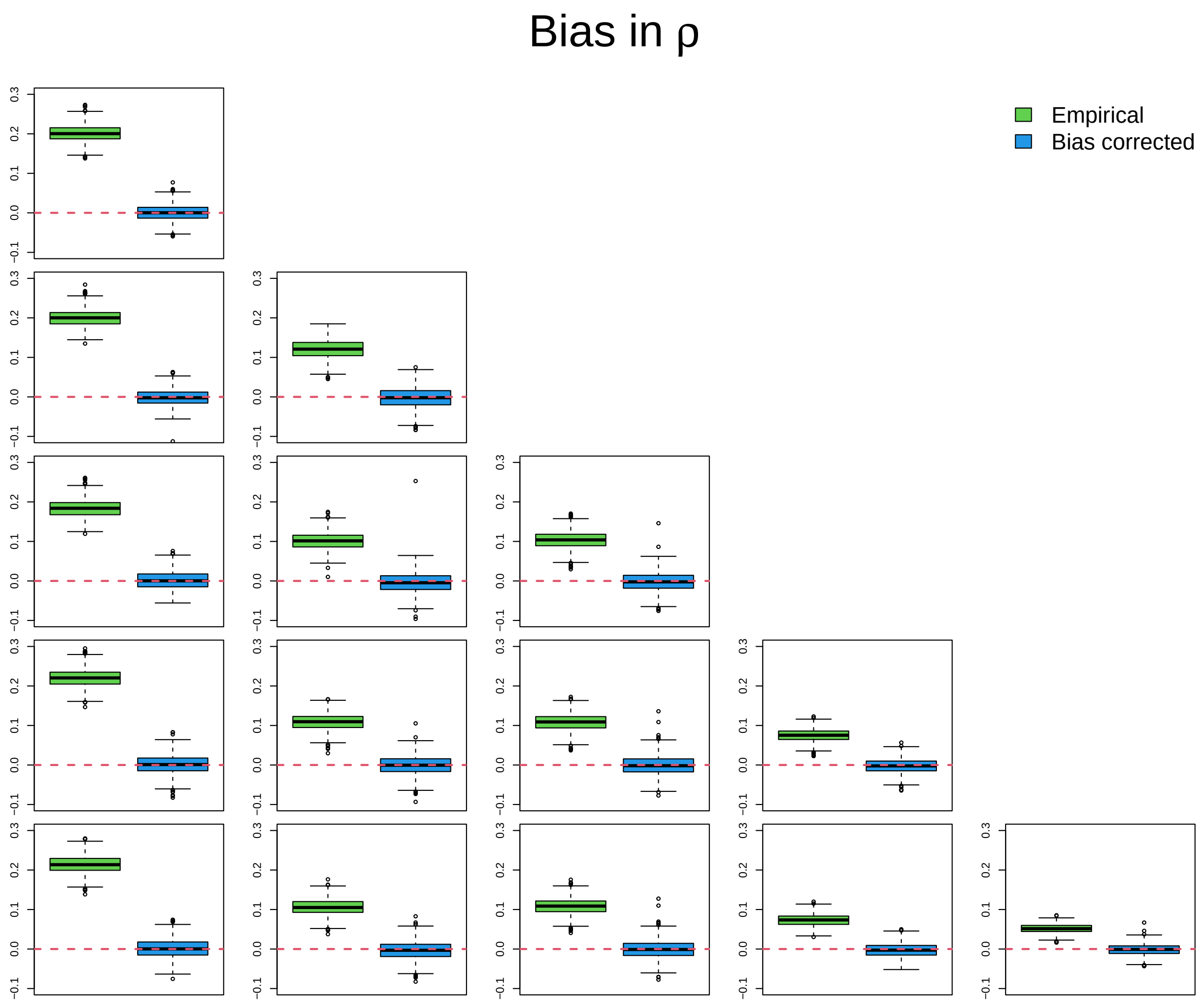


Figure S3: Bias in $\rho(y_{ij}, y_{ik})$ over 1000 simulation replications. The vertical axis is the estimated value minus the true value, $\rho(p_{ij}, p_{ik})$. Each plot corresponds to a pair $(j, k)$ with $j \neq k$. The green boxplots show the empirical estimates, and the blue boxplots are the empirical estimates adjusted using Equation 18.

#### 4 Model fitting procedure

Here we outline the graphical-lasso penalized procedure to fit the logit-normal multinomial model as posited in the main manuscript. This procedure follows Hoff (2003) and Tian (2020). The penalized log-likelihood is given by:

$$\ell\left(\boldsymbol{\mu}, \boldsymbol{\Sigma}^{-1}, \mathbf{w}_1, \ldots, \mathbf{w}_n, \mu_\ell, \sigma_\ell^2\right) \propto \sum_{i=1}^{n}\left[\sum_{j=1}^{J} y_{ij} w_{ij} - n_i \log\left(\sum_{j=1}^{J} e^{w_{ij}} + 1\right)\right]$$
$$+ \frac{n}{2}\log\det\boldsymbol{\Sigma}^{-1} - \frac{1}{2}\sum_{i=1}^{n}(\mathbf{w}_i - \boldsymbol{\mu})^\top\boldsymbol{\Sigma}^{-1}(\mathbf{w}_i - \boldsymbol{\mu}) - \lambda\|\boldsymbol{\Sigma}^{-1}\| \qquad (19)$$
$$- \log(\sigma_\ell) - \frac{1}{2\sigma_\ell^2}\sum_{i=1}^{n}\Big(\log(n_i) - \mu_\ell\Big)^2$$

The penalized maximum likelihood estimates corresponding to Equation 19 can be obtained through an EM procedure. The $\mathbf{w}_i$ parameters are updated using the Newton-Raphson algorithm, where the gradient and Hessian are given by:

$$\nabla\ell(\mathbf{w}_i) = y_i^* - n_i\widehat{\mathbf{p}}_i - \boldsymbol{\Sigma}^{-1}(\mathbf{w}_i - \boldsymbol{\mu})$$
$$\nabla^2\ell(\mathbf{w}_i) = n_i\left(\widehat{\mathbf{p}}_i\widehat{\mathbf{p}}_i^\top - \operatorname{diag}\left(\widehat{\mathbf{p}}_i\right)\right) - \boldsymbol{\Sigma}^{-1},$$

where:

$$\widehat{\mathbf{p}}_i = \frac{\exp(\mathbf{w}_i)}{1 + \sum_{j=1}^{J}\exp(w_{ij})},$$

where $y_i^* = (y_{i1}, \ldots, y_{iJ})$ (i.e. without feature $J+1$), and where the exponential applies element-wise to the vector $\mathbf{w}_i$.

Begin by initializing $\boldsymbol{\mu}^{(0)}$, $(\boldsymbol{\Sigma}^{-1})^{(0)}$, $\mu_\ell^{(0)}$, $\sigma_\ell^{(0)}$. Initialize $\mathbf{w}_1^{(0)}, \ldots, \mathbf{w}_n^{(0)}$ to be the ALR-transformed counts, after imputing zeros, if necessary. Each iteration $k$ of the algorithm then proceeds as follows:

1. Update $\boldsymbol{\mu}^{(k)} = \frac{1}{n}\sum_{i=1}^{n}\mathbf{w}_i^{(k-1)}$
2. Update $(\boldsymbol{\Sigma}^{-1})^{(k)}$ by running graphical lasso on $\mathbf{w}_1^{(k-1)}, \ldots, \mathbf{w}_n^{(k-1)}$.
3. Update $\mathbf{w}_i$ using a Newton-Raphson update with $\nabla\ell(\mathbf{w}_i)$ and $\nabla^2\ell(\mathbf{w}_i)$ defined above.

Finally, the read-depth parameters $\mu_\ell$ and $\sigma_\ell$ are estimated as:

$$\widehat{\mu}_\ell = \frac{1}{n}\sum_{i=1}^{n}\log(n_i) \qquad \widehat{\sigma}_\ell = \frac{1}{n}\sum_{i=1}^{n}\left(\log(n_i) - \widehat{\mu}_\ell\right)^2.$$

To choose the value of the penalization parameter $\lambda$, we fit the model on a range of $\lambda$ values and choose the value that results in the smallest extended Bayesian information criterion (EBIC), which is defined in Chen and Chen (2012).

### 5 Simulation procedure

The simulation parameters were based on an initial multinomial logit-normal model fitted to single-cell RNA-sequencing data from Buettner et al. (2015). In this model, we obtained estimates of $\boldsymbol{\mu}$, $\boldsymbol{\Sigma}$, $\mu_\ell$, and $\sigma_\ell$. To get an estimate of $\boldsymbol{\Omega}$, we apply the second-order approximation as defined in Equation (12) in the main text. The estimated $\boldsymbol{\Omega}$ is then used as the true value of the covariance matrix of the log-proportions in the simulations.

For the sparse $\boldsymbol{\Omega}$ simulation, we choose three sparsity probabilities: $p_{\text{sparse}} \in \{0.1, 0.5, 0.8\}$. For each sparsity probability $p_{\text{sparse}}$, we set the lower $p_{\text{sparse}}$-quantiles of the off-diagonal elements of $\boldsymbol{\Omega}$ to zero. The resulting matrix is positive-definite, so no further adjustment is needed. We then calculate the true value of $\boldsymbol{\Sigma}$ as:

$$\boldsymbol{\Sigma} = \mathbf{F}\boldsymbol{\Omega}\mathbf{F}^\top,$$

$\mathbf{F} = [\mathbf{I}_J, -\mathbf{1}_J]$, with $\mathbf{I}_J$ representing the $J \times J$ identity matrix, and $\mathbf{1}_J$ representing a $J$-dimensional vector of ones. The true value of $\boldsymbol{\mu}$ is used as the estimated value from the data. The simulation proceeds as follows for each $i = 1, \ldots, n$:

1. Generate $\mathbf{w}_i \sim \text{MVN}(\boldsymbol{\mu}, \boldsymbol{\Sigma})$
2. Calculate $\mathbf{p}_i$ as $\text{alr}^{-1}(\mathbf{w}_i)$
3. Simulate $n_i \sim \text{logNormal}(\mu_\ell, \sigma_\ell^2)$
4. Simulate $\mathbf{Y}_i \sim \text{Multinomial}(n_i, \mathbf{p}_i)$

To generate the misspecified model simulation data, we simulated $\mathbf{w}_i$ from a multivariate non-central $t$-distribution with mean and variance equal to $\boldsymbol{\mu}$ and $\boldsymbol{\Sigma}$, and with degrees of freedom fixed to 2.1. All other steps of the simulation procedure remained the same as in the Gaussian version of the simulation. This allows the distribution of latent values $\mathbf{w}_i$ to have heavier tails and be non-symmetric.

### 6 Additional simulation results

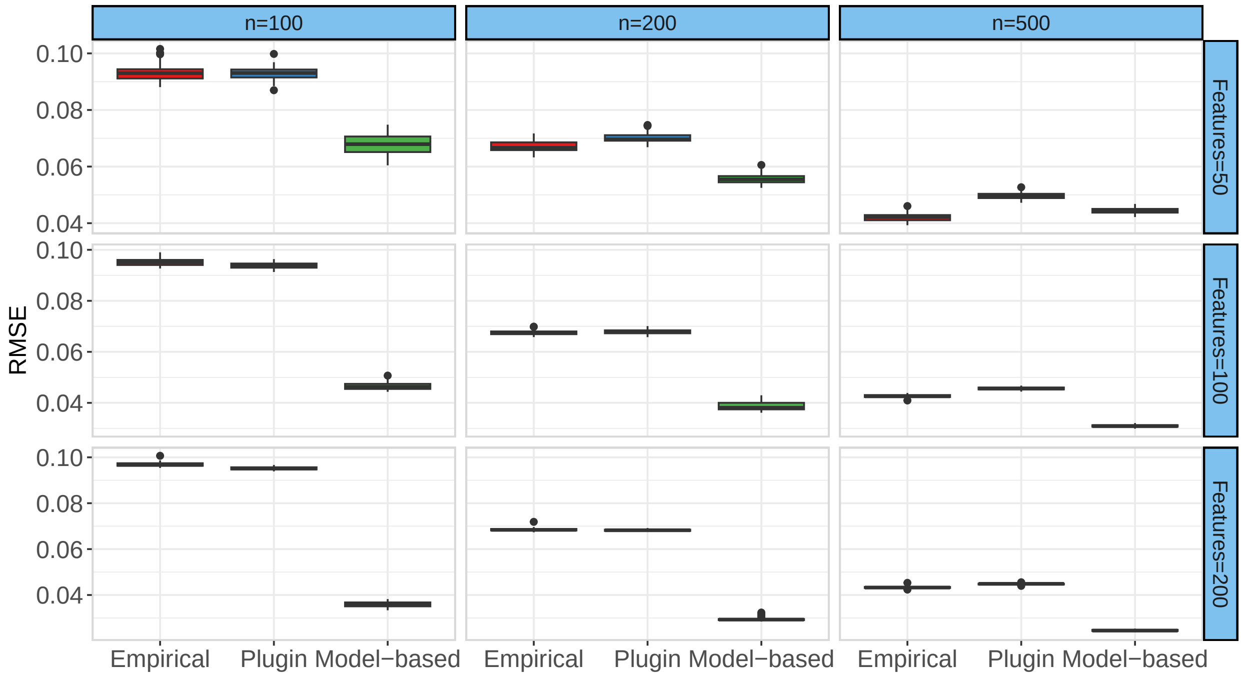


Figure S4: RMSE of $\rho$ estimated empirically and using the model-based procedure. Boxplots are over 50 simulation replications.

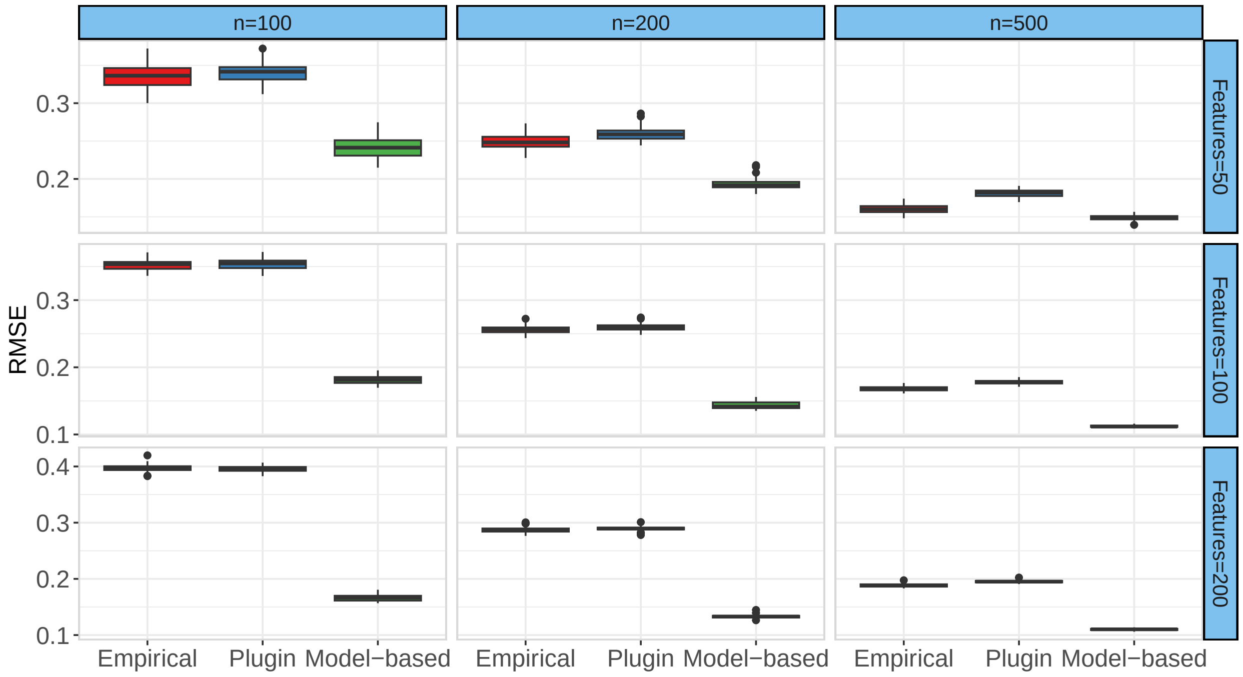


Figure S5: RMSE of $\mathbf{\Omega}$ elements estimated empirically and using the model-based procedure. Boxplots are over 50 simulation replications.

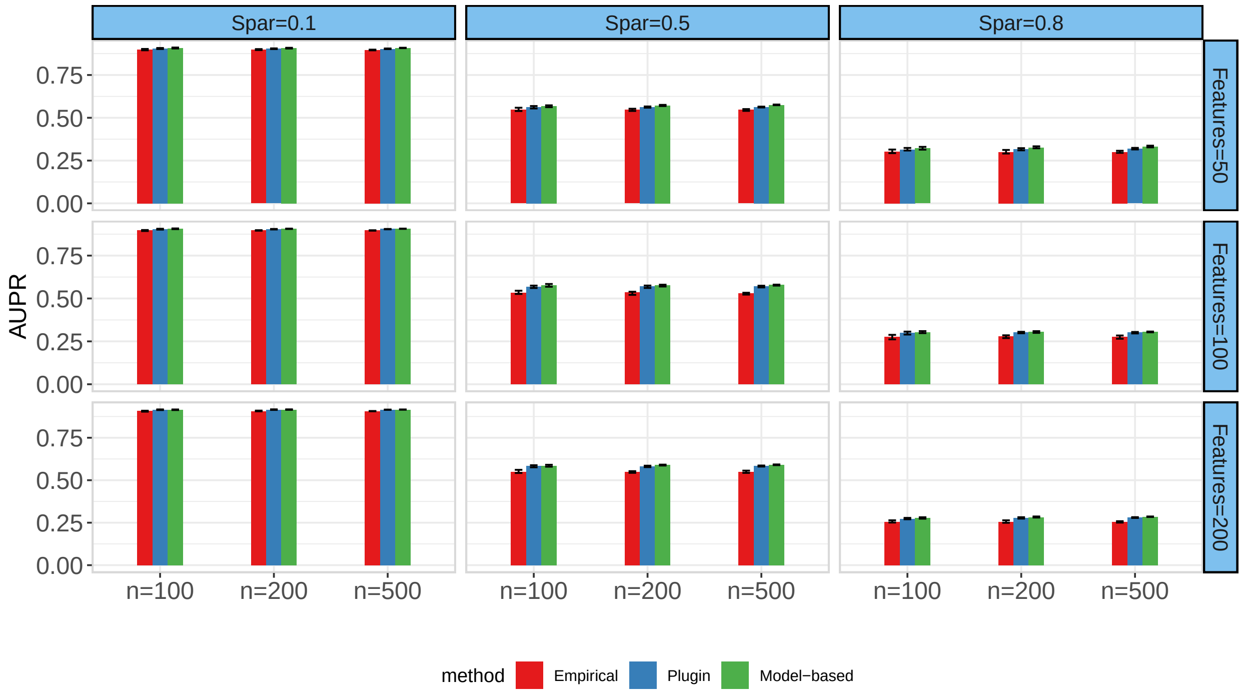


Figure S6: Area under the precision-recall curve of $\rho$ estimation methods in the misspecified model simulation data. Error bars represent 25th and 75th percentiles over 50 simulation replications. Simulations vary over sparsity level (e.g. proportion of true $\rho$ values equal to 0) and number of features $J+1$.

| ID 1 | Assoc. gene 1 | ID 2 | Assoc. gene 2 | $\rho_{G1}$ | $\rho_{G2M}$ | $\rho_{G1} - \rho_{G2M}$ |
|---|---|---|---|---|---|---|
| ENSMUSG00000039187 | Fanci | ENSMUSG00000033417 | Cacul1 | 0.0394 | -0.2453 | 0.2847 |
| ENSMUSG00000064302 | Clasp1 | ENSMUSG00000059248 | Sep-09 | 0.1426 | -0.139 | 0.2816 |
| ENSMUSG00000038482 | Tfdp1 | ENSMUSG00000038416 | Cdc16 | 0.2141 | -0.0585 | 0.2726 |
| ENSMUSG00000040943 | Tet2 | ENSMUSG00000011831 | Evi5 | 0.2575 | -0.0135 | 0.271 |
| ENSMUSG00000029176 | Anapc4 | ENSMUSG00000000001 | Gnai3 | 0.242 | -0.0258 | 0.2678 |
| ENSMUSG00000038619 | Ensa | ENSMUSG00000035683 | Melk | 0.2088 | -0.0562 | 0.2651 |
| ENSMUSG00000079555 | Haus3 | ENSMUSG00000038416 | Cdc16 | 0.0278 | -0.2355 | 0.2633 |
| ENSMUSG00000034023 | Fancd2 | ENSMUSG00000027715 | Ccna2 | 0.1562 | -0.106 | 0.2622 |
| ENSMUSG00000046591 | Ticrr | ENSMUSG00000032113 | Chek1 | 0.0834 | -0.1786 | 0.262 |
| ENSMUSG00000036777 | Anln | ENSMUSG00000021548 | Ccnh | 0.1834 | -0.0779 | 0.2613 |
| ENSMUSG00000036782 | Klhl13 | ENSMUSG00000025616 | Usp16 | 0.0335 | -0.2221 | 0.2556 |
| ENSMUSG00000012443 | Kif11 | ENSMUSG00000010342 | Tex14 | 0.1434 | -0.1074 | 0.2508 |
| ENSMUSG00000039994 | Timeless | ENSMUSG00000019942 | Cdk1 | 0.1917 | -0.0578 | 0.2495 |
| ENSMUSG00000032264 | Zw10 | ENSMUSG00000002068 | Ccne1 | 0.0839 | -0.1638 | 0.2477 |
| ENSMUSG00000061607 | Mdc1 | ENSMUSG00000026779 | Mastl | 0.0184 | -0.228 | 0.2463 |
| ENSMUSG00000035683 | Melk | ENSMUSG00000002297 | Dbf4 | 0.2385 | -0.0073 | 0.2457 |
| ENSMUSG00000032113 | Chek1 | ENSMUSG00000000028 | Cdc45 | 0.2182 | -0.0274 | 0.2456 |
| ENSMUSG00000034023 | Fancd2 | ENSMUSG00000000902 | Smarcb1 | 0.1621 | -0.0807 | 0.2428 |
| ENSMUSG00000025862 | Stag2 | ENSMUSG00000020415 | Pttg1 | 0.1668 | -0.0754 | 0.2422 |
| ENSMUSG00000035683 | Melk | ENSMUSG00000035439 | Haus8 | 0.2334 | -0.0086 | 0.2419 |
| ENSMUSG00000034165 | Ccnd3 | ENSMUSG00000027715 | Ccna2 | 0.1364 | -0.1042 | 0.2406 |
| ENSMUSG00000046591 | Ticrr | ENSMUSG00000029472 | Anapc5 | 0.2067 | -0.0313 | 0.238 |
| ENSMUSG00000030983 | Bccip | ENSMUSG00000028066 | Pmf1 | 0.1458 | -0.0919 | 0.2377 |
| ENSMUSG00000072980 | Oip5 | ENSMUSG00000032477 | Cdc25a | 0.0785 | -0.1579 | 0.2364 |
| ENSMUSG00000038619 | Ensa | ENSMUSG00000019917 | Sep-10 | 0.2036 | -0.0326 | 0.2361 |
| ENSMUSG00000079555 | Haus3 | ENSMUSG00000061607 | Mdc1 | 0.1858 | -0.0487 | 0.2344 |
| ENSMUSG00000034906 | Ncaph | ENSMUSG00000034023 | Fancd2 | 0.226 | -0.0068 | 0.2328 |
| ENSMUSG00000029414 | Kntc1 | ENSMUSG00000019794 | Katna1 | 0.1731 | -0.0542 | 0.2273 |
| ENSMUSG00000041769 | Ppp2r2d | ENSMUSG00000029283 | Cdc7 | 0.1219 | -0.1037 | 0.2256 |
| ENSMUSG00000031314 | Taf1 | ENSMUSG00000029003 | Mad2l2 | 0.1011 | -0.1217 | 0.2228 |
| ENSMUSG00000030105 | Arl8b | ENSMUSG00000004591 | Pkn2 | 0.1846 | -0.0373 | 0.2219 |
| ENSMUSG00000078652 | Psme3 | ENSMUSG00000000001 | Gnai3 | 0.1747 | -0.0465 | 0.2212 |
| ENSMUSG00000031371 | Haus7 | ENSMUSG00000021965 | Ska3 | -0.0199 | 0.2011 | -0.221 |
| ENSMUSG00000039187 | Fanci | ENSMUSG00000025616 | Usp16 | 0.0103 | 0.2313 | -0.221 |
| ENSMUSG00000028896 | Rcc1 | ENSMUSG00000027285 | Haus2 | -0.1073 | 0.1142 | -0.2215 |
| ENSMUSG00000032264 | Zw10 | ENSMUSG00000026361 | Cdc73 | -0.0211 | 0.2016 | -0.2227 |
| ENSMUSG00000059552 | Trp53 | ENSMUSG00000006728 | Cdk4 | -0.0345 | 0.1887 | -0.2232 |
| ENSMUSG00000036672 | Cenpt | ENSMUSG00000019923 | Zwint | 0.0022 | 0.2261 | -0.2239 |
| ENSMUSG00000042750 | Bex2 | ENSMUSG00000039130 | Zc3hc1 | -0.0268 | 0.1977 | -0.2245 |
| ENSMUSG00000034023 | Fancd2 | ENSMUSG00000017499 | Cdc6 | -0.0429 | 0.1816 | -0.2246 |
| ENSMUSG00000072082 | Ccnf | ENSMUSG00000033364 | Usp37 | -0.0648 | 0.1601 | -0.2248 |
| ENSMUSG00000072980 | Oip5 | ENSMUSG00000033054 | Npat | -0.1851 | 0.0437 | -0.2288 |
| ENSMUSG00000026779 | Mastl | ENSMUSG00000021965 | Ska3 | 0.0856 | 0.3146 | -0.2291 |
| ENSMUSG00000068744 | Psrc1 | ENSMUSG00000037725 | Ckap2 | -0.0098 | 0.2193 | -0.2291 |
| ENSMUSG00000033952 | Aspm | ENSMUSG00000025480 | Syce1 | -0.2346 | -0.0045 | -0.2301 |
| ENSMUSG00000038619 | Ensa | ENSMUSG00000026491 | Ahctf1 | -0.0225 | 0.2077 | -0.2302 |
| ENSMUSG00000022678 | Nde1 | ENSMUSG00000020415 | Pttg1 | -0.0344 | 0.1969 | -0.2313 |
| ENSMUSG00000032411 | Tfdp2 | ENSMUSG00000017499 | Cdc6 | -0.2289 | 0.0038 | -0.2327 |
| ENSMUSG00000034023 | Fancd2 | ENSMUSG00000003824 | Syce2 | 0.0083 | 0.2419 | -0.2336 |
| ENSMUSG00000036782 | Klhl13 | ENSMUSG00000020415 | Pttg1 | -0.1079 | 0.1266 | -0.2345 |
| ENSMUSG00000036779 | Papd5 | ENSMUSG00000019923 | Zwint | -0.0192 | 0.2164 | -0.2356 |
| ENSMUSG00000036928 | Stag3 | ENSMUSG00000028896 | Rcc1 | 0.0021 | 0.2382 | -0.2361 |
| ENSMUSG00000073705 | Apitd1 | ENSMUSG00000041997 | Tlk1 | 0.0016 | 0.2384 | -0.2367 |
| ENSMUSG00000031371 | Haus7 | ENSMUSG00000026039 | Sgol2 | -0.043 | 0.1946 | -0.2375 |
| ENSMUSG00000029283 | Cdc7 | ENSMUSG00000025616 | Usp16 | -0.002 | 0.2356 | -0.2376 |
| ENSMUSG00000034165 | Ccnd3 | ENSMUSG00000031371 | Haus7 | -0.0297 | 0.2096 | -0.2392 |
| ENSMUSG00000036850 | Mrpl41 | ENSMUSG00000036782 | Klhl13 | -0.0227 | 0.2166 | -0.2393 |
| ENSMUSG00000038416 | Cdc16 | ENSMUSG00000034023 | Fancd2 | 0.0047 | 0.244 | -0.2393 |
| ENSMUSG00000069631 | Strada | ENSMUSG00000042688 | Mapk6 | -0.1312 | 0.1106 | -0.2418 |
| ENSMUSG00000061607 | Mdc1 | ENSMUSG00000029176 | Anapc4 | -0.1254 | 0.1173 | -0.2427 |
| ENSMUSG00000029003 | Mad2l2 | ENSMUSG00000019794 | Katna1 | -0.0179 | 0.2261 | -0.2439 |
| ENSMUSG00000073705 | Apitd1 | ENSMUSG00000038393 | Txnip | -0.038 | 0.2067 | -0.2447 |
| ENSMUSG00000038902 | Pogz | ENSMUSG00000020415 | Pttg1 | -0.098 | 0.1471 | -0.2452 |
| ENSMUSG00000033054 | Npat | ENSMUSG00000023572 | Ccndbp1 | -0.1109 | 0.1345 | -0.2454 |
| ENSMUSG00000028790 | Khdrbs1 | ENSMUSG00000026965 | Anapc2 | -0.121 | 0.1255 | -0.2465 |
| ENSMUSG00000042029 | Ncapg2 | ENSMUSG00000027363 | Usp8 | -0.0085 | 0.2382 | -0.2467 |
| ENSMUSG00000039187 | Fanci | ENSMUSG00000026965 | Anapc2 | 0.0263 | 0.2737 | -0.2475 |
| ENSMUSG00000039187 | Fanci | ENSMUSG00000025480 | Syce1 | 0.0073 | 0.255 | -0.2477 |
| ENSMUSG00000035683 | Melk | ENSMUSG00000029910 | Mad2l1 | -0.0013 | 0.2466 | -0.2479 |
| ENSMUSG00000038619 | Ensa | ENSMUSG00000036850 | Mrpl41 | 0.0018 | 0.251 | -0.2492 |
| ENSMUSG00000070923 | Klhl9 | ENSMUSG00000059248 | Sep-09 | -0.1331 | 0.1161 | -0.2493 |
| ENSMUSG00000019923 | Zwint | ENSMUSG00000018821 | Avpi1 | -0.004 | 0.246 | -0.25 |
| ENSMUSG00000020415 | Pttg1 | ENSMUSG00000014859 | E2f4 | -0.0806 | 0.1695 | -0.2501 |
| ENSMUSG00000036782 | Klhl13 | ENSMUSG00000021276 | Cinp | -0.0277 | 0.2237 | -0.2514 |
| ENSMUSG00000074476 | Spc24 | ENSMUSG00000019923 | Zwint | -0.0373 | 0.2158 | -0.253 |
| ENSMUSG00000022945 | Chaf1b | ENSMUSG00000019794 | Katna1 | -0.1941 | 0.0594 | -0.2536 |
| ENSMUSG00000036928 | Stag3 | ENSMUSG00000035828 | Pim3 | 0.0128 | 0.2664 | -0.2536 |
| ENSMUSG00000026779 | Mastl | ENSMUSG00000025480 | Syce1 | -0.0276 | 0.2272 | -0.2547 |
| ENSMUSG00000038416 | Cdc16 | ENSMUSG00000018509 | Cenpv | -0.0094 | 0.248 | -0.2574 |
| ENSMUSG00000020415 | Pttg1 | ENSMUSG00000006398 | Cdc20 | -0.0178 | 0.2398 | -0.2577 |
| ENSMUSG00000035683 | Melk | ENSMUSG00000020415 | Pttg1 | -0.1615 | 0.097 | -0.2585 |
| ENSMUSG00000053119 | Chmp3 | ENSMUSG00000022978 | Mis18a | -0.0213 | 0.2377 | -0.259 |
| ENSMUSG00000033054 | Npat | ENSMUSG00000021965 | Ska3 | -0.0064 | 0.2556 | -0.262 |
| ENSMUSG00000038619 | Ensa | ENSMUSG00000036782 | Klhl13 | 0.0052 | 0.2715 | -0.2663 |
| ENSMUSG00000032562 | Gnai2 | ENSMUSG00000027715 | Ccna2 | 0.0037 | 0.274 | -0.2703 |
| ENSMUSG00000042029 | Ncapg2 | ENSMUSG00000038393 | Txnip | -0.1767 | 0.0974 | -0.2741 |
| ENSMUSG00000004085 | B230120H23Rik | ENSMUSG00000002068 | Ccne1 | -0.1921 | 0.0856 | -0.2777 |
| ENSMUSG00000036928 | Stag3 | ENSMUSG00000029176 | Anapc4 | -0.1492 | 0.1296 | -0.2788 |
| ENSMUSG00000035828 | Pim3 | ENSMUSG00000031095 | Cul4b | 0.0019 | 0.2836 | -0.2818 |
| ENSMUSG00000036782 | Klhl13 | ENSMUSG00000021965 | Ska3 | -0.0729 | 0.2097 | -0.2827 |
| ENSMUSG00000038902 | Pogz | ENSMUSG00000031347 | Cetn2 | -0.1185 | 0.1715 | -0.29 |
| ENSMUSG00000023572 | Ccndbp1 | ENSMUSG00000018821 | Avpi1 | 0.0052 | 0.2975 | -0.2923 |
| ENSMUSG00000069631 | Strada | ENSMUSG00000000001 | Gnai3 | -0.1641 | 0.1631 | -0.3271 |
| ENSMUSG00000042688 | Mapk6 | ENSMUSG00000022945 | Chaf1b | -0.2119 | 0.124 | -0.3359 |
| ENSMUSG00000036928 | Stag3 | ENSMUSG00000031371 | Haus7 | -0.0422 | 0.3023 | -0.3445 |
| ENSMUSG00000031371 | Haus7 | ENSMUSG00000026965 | Anapc2 | -0.0151 | 0.3333 | -0.3484 |
| ENSMUSG00000066357 | Wdr6 | ENSMUSG00000021635 | Rad17 | -0.22 | 0.1302 | -0.3502 |
| ENSMUSG00000038619 | Ensa | ENSMUSG00000029176 | Anapc4 | 0.0228 | 0.3732 | -0.3504 |
| ENSMUSG00000073705 | Apitd1 | ENSMUSG00000036928 | Stag3 | 0.0103 | 0.3711 | -0.3608 |
| ENSMUSG00000036928 | Stag3 | ENSMUSG00000021276 | Cinp | -0.0812 | 0.4194 | -0.5005 |

Table S1: Top gene pair differences between G1 and G2M stages.

| ID 1 | Assoc. gene 1 | ID 2 | Assoc. gene 2 | $\rho_{G1}$ | $\rho_S$ | $\rho_{G1} - \rho_S$ |
|---|---|---|---|---|---|---|
| ENSMUSG00000032504 | Pdcd6ip | ENSMUSG00000002635 | Pdcd2l | **0.0057** | **0.4444** | **-0.4387** |
| ENSMUSG00000042029 | Ncapg2 | ENSMUSG00000022945 | Chaf1b | **-0.0163** | **0.4252** | **-0.4415** |
| ENSMUSG00000034218 | Atm | ENSMUSG00000000001 | Gnai3 | **-0.0296** | **0.4123** | **-0.4419** |
| ENSMUSG00000046591 | Ticrr | ENSMUSG00000000708 | Kat2b | **-0.1192** | **0.3236** | **-0.4428** |
| ENSMUSG00000036782 | Klhl13 | ENSMUSG00000029521 | Chek2 | **-0.0181** | **0.4252** | **-0.4433** |
| ENSMUSG00000038393 | Txnip | ENSMUSG00000020107 | Anapc16 | **0.0241** | **0.4681** | **-0.4439** |
| ENSMUSG00000057110 | Cep110 | ENSMUSG00000034218 | Atm | **0.0597** | **0.5042** | **-0.4445** |
| ENSMUSG00000039128 | Cdc123 | ENSMUSG00000033417 | Cacul1 | **-0.0113** | **0.434** | **-0.4453** |
| ENSMUSG00000027550 | Lrrcc1 | ENSMUSG00000000001 | Gnai3 | **-0.0563** | **0.3896** | **-0.4459** |
| ENSMUSG00000032400 | Zwilch | ENSMUSG00000029176 | Anapc4 | **-0.0346** | **0.4132** | **-0.4478** |
| ENSMUSG00000045969 | Ing1 | ENSMUSG00000031858 | Mau2 | **0.01** | **0.4592** | **-0.4491** |
| ENSMUSG00000029501 | Ankle2 | ENSMUSG00000028873 | Cdca8 | **0.009** | **0.4596** | **-0.4507** |
| ENSMUSG00000014859 | E2f4 | ENSMUSG00000000001 | Gnai3 | **0.0218** | **0.4732** | **-0.4514** |
| ENSMUSG00000045328 | Cenpe | ENSMUSG00000017146 | Brca1 | **-0.0055** | **0.4462** | **-0.4517** |
| ENSMUSG00000033054 | Npat | ENSMUSG00000022945 | Chaf1b | **-0.0626** | **0.3894** | **-0.452** |
| ENSMUSG00000026779 | Mastl | ENSMUSG00000010342 | Tex14 | **0.0471** | **0.5003** | **-0.4533** |
| ENSMUSG00000033054 | Npat | ENSMUSG00000029501 | Ankle2 | **0.0065** | **0.4597** | **-0.4533** |
| ENSMUSG00000029003 | Mad2l2 | ENSMUSG00000023572 | Ccndbp1 | **0.0038** | **0.4579** | **-0.4541** |
| ENSMUSG00000069631 | Strada | ENSMUSG00000000001 | Gnai3 | **-0.1641** | **0.2905** | **-0.4546** |
| ENSMUSG00000066149 | Cdc26 | ENSMUSG00000038393 | Txnip | **-0.0172** | **0.4377** | **-0.455** |
| ENSMUSG00000031858 | Mau2 | ENSMUSG00000029176 | Anapc4 | **0.0567** | **0.5129** | **-0.4561** |
| ENSMUSG00000032113 | Chek1 | ENSMUSG00000020107 | Anapc16 | **-0.0292** | **0.4269** | **-0.4561** |
| ENSMUSG00000034218 | Atm | ENSMUSG00000030641 | 4632434I11Rik | **-0.008** | **0.448** | **-0.4561** |
| ENSMUSG00000032113 | Chek1 | ENSMUSG00000029003 | Mad2l2 | **0.0307** | **0.4883** | **-0.4576** |
| ENSMUSG00000032113 | Chek1 | ENSMUSG00000024943 | Smc5 | **-0.0193** | **0.4403** | **-0.4596** |
| ENSMUSG00000030641 | 4632434I11Rik | ENSMUSG00000021595 | Nsun2 | **0.0198** | **0.48** | **-0.4602** |
| ENSMUSG00000033417 | Cacul1 | ENSMUSG00000000001 | Gnai3 | **-0.0106** | **0.4501** | **-0.4607** |
| ENSMUSG00000041238 | Rbbp8 | ENSMUSG00000032113 | Chek1 | **-0.0176** | **0.4462** | **-0.4638** |
| ENSMUSG00000038252 | Ncapd2 | ENSMUSG00000020694 | Tlk2 | **-0.0051** | **0.4592** | **-0.4643** |
| ENSMUSG00000029147 | Ppm1g | ENSMUSG00000028059 | Arhgef2 | **0.0132** | **0.4782** | **-0.465** |
| ENSMUSG00000045969 | Ing1 | ENSMUSG00000029176 | Anapc4 | **0.0293** | **0.495** | **-0.4657** |
| ENSMUSG00000057110 | Cep110 | ENSMUSG00000026779 | Mastl | **0.0541** | **0.5198** | **-0.4657** |
| ENSMUSG00000031371 | Haus7 | ENSMUSG00000020107 | Anapc16 | **-0.0514** | **0.4146** | **-0.4659** |
| ENSMUSG00000041238 | Rbbp8 | ENSMUSG00000038393 | Txnip | **-0.0119** | **0.4539** | **-0.4659** |
| ENSMUSG00000029253 | Cenpc1 | ENSMUSG00000020694 | Tlk2 | **0.026** | **0.492** | **-0.466** |
| ENSMUSG00000030061 | Uba3 | ENSMUSG00000029501 | Ankle2 | **0.0046** | **0.4706** | **-0.4661** |
| ENSMUSG00000045328 | Cenpe | ENSMUSG00000028678 | Kif2c | **0.042** | **0.5106** | **-0.4686** |
| ENSMUSG00000041238 | Rbbp8 | ENSMUSG00000030641 | 4632434I11Rik | **-0.0179** | **0.4517** | **-0.4695** |
| ENSMUSG00000034218 | Atm | ENSMUSG00000029501 | Ankle2 | **0.0012** | **0.4715** | **-0.4703** |
| ENSMUSG00000038619 | Ensa | ENSMUSG00000032113 | Chek1 | **0.0126** | **0.484** | **-0.4715** |
| ENSMUSG00000032113 | Chek1 | ENSMUSG00000025862 | Stag2 | **0.0962** | **0.5679** | **-0.4717** |
| ENSMUSG00000034218 | Atm | ENSMUSG00000022945 | Chaf1b | **0.0202** | **0.4934** | **-0.4732** |
| ENSMUSG00000014859 | E2f4 | ENSMUSG00000010342 | Tex14 | **-0.0622** | **0.4122** | **-0.4744** |
| ENSMUSG00000029501 | Ankle2 | ENSMUSG00000029003 | Mad2l2 | **0.0042** | **0.4787** | **-0.4745** |
| ENSMUSG00000034023 | Fancd2 | ENSMUSG00000011831 | Evi5 | **-0.1067** | **0.368** | **-0.4747** |
| ENSMUSG00000032113 | Chek1 | ENSMUSG00000022945 | Chaf1b | **0.0455** | **0.5203** | **-0.4748** |
| ENSMUSG00000038393 | Txnip | ENSMUSG00000030061 | Uba3 | **0.0311** | **0.5072** | **-0.4761** |
| ENSMUSG00000066149 | Cdc26 | ENSMUSG00000020107 | Anapc16 | **0.0389** | **0.5152** | **-0.4764** |
| ENSMUSG00000032113 | Chek1 | ENSMUSG00000029147 | Ppm1g | **-0.0713** | **0.4063** | **-0.4776** |
| ENSMUSG00000061665 | Cd2ap | ENSMUSG00000041238 | Rbbp8 | **-0.1598** | **0.3182** | **-0.478** |
| ENSMUSG00000031858 | Mau2 | ENSMUSG00000029147 | Ppm1g | **0.0133** | **0.4934** | **-0.4801** |
| ENSMUSG00000029501 | Ankle2 | ENSMUSG00000014355 | Anapc1 | **0.02** | **0.501** | **-0.481** |
| ENSMUSG00000036782 | Klhl13 | ENSMUSG00000034218 | Atm | **-0.0326** | **0.4484** | **-0.481** |
| ENSMUSG00000030641 | 4632434I11Rik | ENSMUSG00000023572 | Ccndbp1 | **-0.0287** | **0.4537** | **-0.4824** |
| ENSMUSG00000032113 | Chek1 | ENSMUSG00000014355 | Anapc1 | **0.0046** | **0.4897** | **-0.4851** |
| ENSMUSG00000034218 | Atm | ENSMUSG00000031371 | Haus7 | **0.0081** | **0.4969** | **-0.4888** |
| ENSMUSG00000029501 | Ankle2 | ENSMUSG00000022945 | Chaf1b | **-0.0145** | **0.4761** | **-0.4906** |
| ENSMUSG00000057110 | Cep110 | ENSMUSG00000014859 | E2f4 | **0.0034** | **0.4946** | **-0.4912** |
| ENSMUSG00000030649 | 3200002M19Rik | ENSMUSG00000010342 | Tex14 | **0.034** | **0.5284** | **-0.4944** |
| ENSMUSG00000069631 | Strada | ENSMUSG00000045969 | Ing1 | **-0.0097** | **0.489** | **-0.4987** |
| ENSMUSG00000029147 | Ppm1g | ENSMUSG00000025862 | Stag2 | **-0.0251** | **0.475** | **-0.5001** |
| ENSMUSG00000066149 | Cdc26 | ENSMUSG00000020694 | Tlk2 | **0.0074** | **0.5087** | **-0.5013** |
| ENSMUSG00000032113 | Chek1 | ENSMUSG00000030641 | 4632434I11Rik | **0.0127** | **0.517** | **-0.5043** |
| ENSMUSG00000029176 | Anapc4 | ENSMUSG00000029147 | Ppm1g | **0.023** | **0.5288** | **-0.5058** |
| ENSMUSG00000033054 | Npat | ENSMUSG00000025862 | Stag2 | **-0.0767** | **0.4336** | **-0.5102** |
| ENSMUSG00000034218 | Atm | ENSMUSG00000029176 | Anapc4 | **-0.0047** | **0.5059** | **-0.5106** |
| ENSMUSG00000029253 | Cenpc1 | ENSMUSG00000000708 | Kat2b | **-0.0038** | **0.5073** | **-0.5111** |
| ENSMUSG00000045969 | Ing1 | ENSMUSG00000032113 | Chek1 | **-0.0061** | **0.506** | **-0.5121** |
| ENSMUSG00000030641 | 4632434I11Rik | ENSMUSG00000029003 | Mad2l2 | **-0.0059** | **0.5089** | **-0.5148** |
| ENSMUSG00000014859 | E2f4 | ENSMUSG00000000708 | Kat2b | **-0.0297** | **0.4876** | **-0.5174** |
| ENSMUSG00000034218 | Atm | ENSMUSG00000029003 | Mad2l2 | **-0.0685** | **0.4523** | **-0.5208** |
| ENSMUSG00000038393 | Txnip | ENSMUSG00000032113 | Chek1 | **0.0233** | **0.5449** | **-0.5216** |
| ENSMUSG00000038393 | Txnip | ENSMUSG00000029501 | Ankle2 | **-0.0117** | **0.5114** | **-0.5231** |
| ENSMUSG00000041238 | Rbbp8 | ENSMUSG00000029501 | Ankle2 | **0.0064** | **0.5325** | **-0.5262** |
| ENSMUSG00000033054 | Npat | ENSMUSG00000029176 | Anapc4 | **-0.0297** | **0.498** | **-0.5277** |
| ENSMUSG00000033054 | Npat | ENSMUSG00000031858 | Mau2 | **0.0186** | **0.5476** | **-0.5289** |
| ENSMUSG00000022945 | Chaf1b | ENSMUSG00000019794 | Katna1 | **-0.1941** | **0.3376** | **-0.5318** |
| ENSMUSG00000045969 | Ing1 | ENSMUSG00000028059 | Arhgef2 | **0.0212** | **0.5558** | **-0.5346** |
| ENSMUSG00000057110 | Cep110 | ENSMUSG00000010342 | Tex14 | **-0.0248** | **0.5168** | **-0.5416** |
| ENSMUSG00000066149 | Cdc26 | ENSMUSG00000029253 | Cenpc1 | **0.0308** | **0.5738** | **-0.543** |
| ENSMUSG00000057110 | Cep110 | ENSMUSG00000000001 | Gnai3 | **0.0063** | **0.5515** | **-0.5452** |
| ENSMUSG00000029176 | Anapc4 | ENSMUSG00000025862 | Stag2 | **0.0287** | **0.5803** | **-0.5516** |
| ENSMUSG00000034218 | Atm | ENSMUSG00000033054 | Npat | **-0.1017** | **0.4514** | **-0.5531** |
| ENSMUSG00000031858 | Mau2 | ENSMUSG00000028059 | Arhgef2 | **0.0228** | **0.5783** | **-0.5555** |
| ENSMUSG00000031371 | Haus7 | ENSMUSG00000029501 | Ankle2 | **-0.0339** | **0.5222** | **-0.5562** |
| ENSMUSG00000066149 | Cdc26 | ENSMUSG00000000708 | Kat2b | **0.0144** | **0.5754** | **-0.5609** |
| ENSMUSG00000031858 | Mau2 | ENSMUSG00000025862 | Stag2 | **-0.0235** | **0.5384** | **-0.5619** |
| ENSMUSG00000033054 | Npat | ENSMUSG00000028059 | Arhgef2 | **0.0238** | **0.5868** | **-0.5629** |
| ENSMUSG00000034218 | Atm | ENSMUSG00000010342 | Tex14 | **-0.0147** | **0.5573** | **-0.572** |
| ENSMUSG00000034218 | Atm | ENSMUSG00000032113 | Chek1 | **0.0074** | **0.5869** | **-0.5796** |
| ENSMUSG00000032113 | Chek1 | ENSMUSG00000029501 | Ankle2 | **-0.0151** | **0.5749** | **-0.59** |
| ENSMUSG00000034218 | Atm | ENSMUSG00000026779 | Mastl | **0.0037** | **0.6005** | **-0.5969** |
| ENSMUSG00000032113 | Chek1 | ENSMUSG00000029176 | Anapc4 | **-0.0744** | **0.5354** | **-0.6098** |
| ENSMUSG00000033054 | Npat | ENSMUSG00000032113 | Chek1 | **0.01** | **0.6316** | **-0.6216** |
| ENSMUSG00000032113 | Chek1 | ENSMUSG00000023572 | Ccndbp1 | **-0.1321** | **0.4908** | **-0.6229** |
| ENSMUSG00000032113 | Chek1 | ENSMUSG00000031858 | Mau2 | **1e-04** | **0.6293** | **-0.6292** |
| ENSMUSG00000029176 | Anapc4 | ENSMUSG00000028059 | Arhgef2 | **-0.0212** | **0.6268** | **-0.6479** |
| ENSMUSG00000032113 | Chek1 | ENSMUSG00000028059 | Arhgef2 | **0.0159** | **0.6667** | **-0.6508** |
| ENSMUSG00000028059 | Arhgef2 | ENSMUSG00000025862 | Stag2 | **0.0349** | **0.6874** | **-0.6525** |
| ENSMUSG00000034218 | Atm | ENSMUSG00000028059 | Arhgef2 | **0.0201** | **0.6812** | **-0.6611** |

Table S2: Top gene pair differences between G1 and S stages.

| ID 1 | Assoc. gene 1 | ID 2 | Assoc. gene 2 | $\rho_{G2M}$ | $\rho_S$ | $\rho_{G2M} - \rho_S$ |
|---|---|---|---|---|---|---|
| ENSMUSG00000022945 | Chaf1b | ENSMUSG00000010342 | Tex14 | -0.0537 | 0.3845 | -0.4382 |
| ENSMUSG00000034218 | Atm | ENSMUSG00000029003 | Mad2l2 | 0.0135 | 0.4523 | -0.4387 |
| ENSMUSG00000042029 | Ncapg2 | ENSMUSG00000022945 | Chaf1b | -0.0144 | 0.4252 | -0.4396 |
| ENSMUSG00000032113 | Chek1 | ENSMUSG00000031371 | Haus7 | -0.02 | 0.4198 | -0.4399 |
| ENSMUSG00000032113 | Chek1 | ENSMUSG00000030061 | Uba3 | 0.0011 | 0.4411 | -0.44 |
| ENSMUSG00000038393 | Txnip | ENSMUSG00000034023 | Fancd2 | -0.105 | 0.3352 | -0.4402 |
| ENSMUSG00000034218 | Atm | ENSMUSG00000029501 | Ankle2 | 0.0311 | 0.4715 | -0.4405 |
| ENSMUSG00000069631 | Strada | ENSMUSG00000039128 | Cdc123 | -0.0215 | 0.4196 | -0.4412 |
| ENSMUSG00000032113 | Chek1 | ENSMUSG00000031858 | Mau2 | 0.1878 | 0.6293 | -0.4415 |
| ENSMUSG00000033054 | Npat | ENSMUSG00000029501 | Ankle2 | 0.0174 | 0.4597 | -0.4424 |
| ENSMUSG00000078652 | Psme3 | ENSMUSG00000029003 | Mad2l2 | -0.0843 | 0.3585 | -0.4428 |
| ENSMUSG00000005410 | Mcm5 | ENSMUSG00000004451 | Ralb | -0.0497 | 0.3939 | -0.4436 |
| ENSMUSG00000061665 | Cd2ap | ENSMUSG00000038393 | Txnip | 0.0091 | 0.4531 | -0.4439 |
| ENSMUSG00000033054 | Npat | ENSMUSG00000029147 | Ppm1g | 0.0092 | 0.4537 | -0.4445 |
| ENSMUSG00000039128 | Cdc123 | ENSMUSG00000023572 | Ccndbp1 | 0.0562 | 0.5008 | -0.4446 |
| ENSMUSG00000032504 | Pdcd6ip | ENSMUSG00000002635 | Pdcd2l | -3e-04 | 0.4444 | -0.4447 |
| ENSMUSG00000029501 | Ankle2 | ENSMUSG00000021965 | Ska3 | -0.0252 | 0.4208 | -0.4459 |
| ENSMUSG00000031858 | Mau2 | ENSMUSG00000029147 | Ppm1g | 0.0473 | 0.4934 | -0.4461 |
| ENSMUSG00000038252 | Ncapd2 | ENSMUSG00000027379 | Bub1 | -0.0329 | 0.4138 | -0.4467 |
| ENSMUSG00000045328 | Cenpe | ENSMUSG00000017146 | Brca1 | -6e-04 | 0.4462 | -0.4468 |
| ENSMUSG00000038619 | Ensa | ENSMUSG00000032113 | Chek1 | 0.0368 | 0.484 | -0.4473 |
| ENSMUSG00000029501 | Ankle2 | ENSMUSG00000020107 | Anapc16 | -9e-04 | 0.4465 | -0.4474 |
| ENSMUSG00000029003 | Mad2l2 | ENSMUSG00000014355 | Anapc1 | -0.0337 | 0.4144 | -0.4481 |
| ENSMUSG00000066149 | Cdc26 | ENSMUSG00000038393 | Txnip | -0.0104 | 0.4377 | -0.4481 |
| ENSMUSG00000033054 | Npat | ENSMUSG00000031858 | Mau2 | 0.0974 | 0.5476 | -0.4501 |
| ENSMUSG00000029003 | Mad2l2 | ENSMUSG00000028678 | Kif2c | -0.1037 | 0.3483 | -0.4521 |
| ENSMUSG00000045969 | Ing1 | ENSMUSG00000032113 | Chek1 | 0.0498 | 0.506 | -0.4562 |
| ENSMUSG00000041238 | Rbbp8 | ENSMUSG00000038393 | Txnip | -0.0041 | 0.4539 | -0.458 |
| ENSMUSG00000045969 | Ing1 | ENSMUSG00000029176 | Anapc4 | 0.0367 | 0.495 | -0.4583 |
| ENSMUSG00000078652 | Psme3 | ENSMUSG00000045328 | Cenpe | -0.0111 | 0.4475 | -0.4586 |
| ENSMUSG00000014859 | E2f4 | ENSMUSG00000000708 | Kat2b | 0.0283 | 0.4876 | -0.4593 |
| ENSMUSG00000057110 | Cep110 | ENSMUSG00000034218 | Atm | 0.0444 | 0.5042 | -0.4598 |
| ENSMUSG00000031858 | Mau2 | ENSMUSG00000029176 | Anapc4 | 0.051 | 0.5129 | -0.4619 |
| ENSMUSG00000055024 | Ep300 | ENSMUSG00000021595 | Nsun2 | -0.0311 | 0.4314 | -0.4626 |
| ENSMUSG00000069631 | Strada | ENSMUSG00000045969 | Ing1 | 0.0255 | 0.489 | -0.4635 |
| ENSMUSG00000003824 | Syce2 | ENSMUSG00000000001 | Gnai3 | -0.0861 | 0.3776 | -0.4637 |
| ENSMUSG00000029253 | Cenpc1 | ENSMUSG00000020694 | Tlk2 | 0.0271 | 0.492 | -0.465 |
| ENSMUSG00000057110 | Cep110 | ENSMUSG00000005410 | Mcm5 | -0.0203 | 0.4447 | -0.465 |
| ENSMUSG00000034218 | Atm | ENSMUSG00000030649 | 3200002M19Rik | 0.0478 | 0.514 | -0.4662 |
| ENSMUSG00000029501 | Ankle2 | ENSMUSG00000029003 | Mad2l2 | 0.012 | 0.4787 | -0.4667 |
| ENSMUSG00000041997 | Tlk1 | ENSMUSG00000025907 | Rb1cc1 | -0.166 | 0.3034 | -0.4694 |
| ENSMUSG00000038393 | Txnip | ENSMUSG00000029147 | Ppm1g | -0.1762 | 0.2943 | -0.4705 |
| ENSMUSG00000033054 | Npat | ENSMUSG00000025862 | Stag2 | -0.0378 | 0.4336 | -0.4714 |
| ENSMUSG00000066149 | Cdc26 | ENSMUSG00000020694 | Tlk2 | 0.0354 | 0.5087 | -0.4732 |
| ENSMUSG00000027550 | Lrrcc1 | ENSMUSG00000023572 | Ccndbp1 | -0.1443 | 0.3291 | -0.4735 |
| ENSMUSG00000032113 | Chek1 | ENSMUSG00000014355 | Anapc1 | 0.0152 | 0.4897 | -0.4745 |
| ENSMUSG00000057110 | Cep110 | ENSMUSG00000014859 | E2f4 | 0.0179 | 0.4946 | -0.4767 |
| ENSMUSG00000028896 | Rcc1 | ENSMUSG00000028059 | Arhgef2 | -0.0687 | 0.4087 | -0.4774 |
| ENSMUSG00000034218 | Atm | ENSMUSG00000029176 | Anapc4 | 0.0281 | 0.5059 | -0.4778 |
| ENSMUSG00000035351 | Nup37 | ENSMUSG00000021595 | Nsun2 | -0.1255 | 0.3527 | -0.4782 |
| ENSMUSG00000032504 | Pdcd6ip | ENSMUSG00000025144 | Stra13 | -0.0418 | 0.4386 | -0.4803 |
| ENSMUSG00000030061 | Uba3 | ENSMUSG00000029501 | Ankle2 | -0.0103 | 0.4706 | -0.4809 |
| ENSMUSG00000040084 | Bub1b | ENSMUSG00000021103 | Mnat1 | -0.0318 | 0.4497 | -0.4815 |
| ENSMUSG00000029253 | Cenpc1 | ENSMUSG00000000708 | Kat2b | 0.0221 | 0.5073 | -0.4853 |
| ENSMUSG00000066149 | Cdc26 | ENSMUSG00000020107 | Anapc16 | 0.0292 | 0.5152 | -0.4861 |
| ENSMUSG00000032113 | Chek1 | ENSMUSG00000030641 | 4632434I11Rik | 0.0263 | 0.517 | -0.4907 |
| ENSMUSG00000045328 | Cenpe | ENSMUSG00000038393 | Txnip | -0.0362 | 0.4552 | -0.4915 |
| ENSMUSG00000045969 | Ing1 | ENSMUSG00000025862 | Stag2 | -0.0144 | 0.481 | -0.4954 |
| ENSMUSG00000032113 | Chek1 | ENSMUSG00000023572 | Ccndbp1 | -0.005 | 0.4908 | -0.4957 |
| ENSMUSG00000029147 | Ppm1g | ENSMUSG00000028059 | Arhgef2 | -0.0178 | 0.4782 | -0.496 |
| ENSMUSG00000032113 | Chek1 | ENSMUSG00000029176 | Anapc4 | 0.0381 | 0.5354 | -0.4973 |
| ENSMUSG00000030641 | 4632434I11Rik | ENSMUSG00000021595 | Nsun2 | -0.0195 | 0.48 | -0.4996 |
| ENSMUSG00000033054 | Npat | ENSMUSG00000014355 | Anapc1 | -0.1327 | 0.3671 | -0.4998 |
| ENSMUSG00000032113 | Chek1 | ENSMUSG00000029501 | Ankle2 | 0.0724 | 0.5749 | -0.5025 |
| ENSMUSG00000026779 | Mastl | ENSMUSG00000010342 | Tex14 | -0.0055 | 0.5003 | -0.5058 |
| ENSMUSG00000034218 | Atm | ENSMUSG00000025862 | Stag2 | -0.0256 | 0.4904 | -0.516 |
| ENSMUSG00000010342 | Tex14 | ENSMUSG00000000001 | Gnai3 | -0.0419 | 0.4757 | -0.5176 |
| ENSMUSG00000029521 | Chek2 | ENSMUSG00000026779 | Mastl | -0.1033 | 0.4178 | -0.5211 |
| ENSMUSG00000029501 | Ankle2 | ENSMUSG00000014355 | Anapc1 | -0.0246 | 0.501 | -0.5256 |
| ENSMUSG00000032113 | Chek1 | ENSMUSG00000029003 | Mad2l2 | -0.0393 | 0.4883 | -0.5276 |
| ENSMUSG00000041238 | Rbbp8 | ENSMUSG00000029003 | Mad2l2 | -0.1168 | 0.4124 | -0.5292 |
| ENSMUSG00000057110 | Cep110 | ENSMUSG00000026779 | Mastl | -0.0096 | 0.5198 | -0.5294 |
| ENSMUSG00000030649 | 3200002M19Rik | ENSMUSG00000000001 | Gnai3 | -0.1098 | 0.4199 | -0.5296 |
| ENSMUSG00000066149 | Cdc26 | ENSMUSG00000029253 | Cenpc1 | 0.0436 | 0.5738 | -0.5301 |
| ENSMUSG00000030649 | 3200002M19Rik | ENSMUSG00000010342 | Tex14 | -0.002 | 0.5284 | -0.5303 |
| ENSMUSG00000029147 | Ppm1g | ENSMUSG00000025862 | Stag2 | -0.0558 | 0.475 | -0.5308 |
| ENSMUSG00000034218 | Atm | ENSMUSG00000031371 | Haus7 | -0.0346 | 0.4969 | -0.5314 |
| ENSMUSG00000041238 | Rbbp8 | ENSMUSG00000029501 | Ankle2 | -0.0011 | 0.5325 | -0.5337 |
| ENSMUSG00000045328 | Cenpe | ENSMUSG00000028678 | Kif2c | -0.0296 | 0.5106 | -0.5402 |
| ENSMUSG00000034218 | Atm | ENSMUSG00000032113 | Chek1 | 0.0439 | 0.5869 | -0.543 |
| ENSMUSG00000029176 | Anapc4 | ENSMUSG00000025862 | Stag2 | 0.0311 | 0.5803 | -0.5492 |
| ENSMUSG00000031371 | Haus7 | ENSMUSG00000029501 | Ankle2 | -0.0293 | 0.5222 | -0.5515 |
| ENSMUSG00000066149 | Cdc26 | ENSMUSG00000000708 | Kat2b | 0.0216 | 0.5754 | -0.5538 |
| ENSMUSG00000057110 | Cep110 | ENSMUSG00000010342 | Tex14 | -0.0371 | 0.5168 | -0.5539 |
| ENSMUSG00000057110 | Cep110 | ENSMUSG00000000001 | Gnai3 | -0.0062 | 0.5515 | -0.5577 |
| ENSMUSG00000031858 | Mau2 | ENSMUSG00000025862 | Stag2 | -0.0207 | 0.5384 | -0.5591 |
| ENSMUSG00000038393 | Txnip | ENSMUSG00000029501 | Ankle2 | -0.0497 | 0.5114 | -0.5611 |
| ENSMUSG00000031858 | Mau2 | ENSMUSG00000028059 | Arhgef2 | 0.008 | 0.5783 | -0.5703 |
| ENSMUSG00000033054 | Npat | ENSMUSG00000032113 | Chek1 | 0.0593 | 0.6316 | -0.5723 |
| ENSMUSG00000033054 | Npat | ENSMUSG00000028059 | Arhgef2 | 0.0144 | 0.5868 | -0.5724 |
| ENSMUSG00000034218 | Atm | ENSMUSG00000010342 | Tex14 | -0.0162 | 0.5573 | -0.5735 |
| ENSMUSG00000045969 | Ing1 | ENSMUSG00000028059 | Arhgef2 | -0.0332 | 0.5558 | -0.589 |
| ENSMUSG00000038393 | Txnip | ENSMUSG00000032113 | Chek1 | -0.0446 | 0.5449 | -0.5894 |
| ENSMUSG00000032113 | Chek1 | ENSMUSG00000025862 | Stag2 | -0.0426 | 0.5679 | -0.6104 |
| ENSMUSG00000034218 | Atm | ENSMUSG00000026779 | Mastl | -0.0148 | 0.6005 | -0.6154 |
| ENSMUSG00000029176 | Anapc4 | ENSMUSG00000028059 | Arhgef2 | 0.003 | 0.6268 | -0.6238 |
| ENSMUSG00000029176 | Anapc4 | ENSMUSG00000029147 | Ppm1g | -0.1249 | 0.5288 | -0.6537 |
| ENSMUSG00000028059 | Arhgef2 | ENSMUSG00000025862 | Stag2 | 0.0101 | 0.6874 | -0.6773 |
| ENSMUSG00000032113 | Chek1 | ENSMUSG00000028059 | Arhgef2 | -0.0147 | 0.6667 | -0.6813 |
| ENSMUSG00000034218 | Atm | ENSMUSG00000028059 | Arhgef2 | -0.0129 | 0.6812 | -0.6941 |

Table S3: Top gene pair differences between G2M and S stages.

| ID 1 | Assoc. gene 1 | ID 2 | Assoc. gene 2 | $\rho_{G1}$ |
|---|---|---|---|---|
| ENSMUSG00000040943 | Tet2 | ENSMUSG00000011831 | Evi5 | **0.2575** |
| ENSMUSG00000049932 | H2afx | ENSMUSG00000025925 | Terf1 | **0.2465** |
| ENSMUSG00000038902 | Pogz | ENSMUSG00000031858 | Mau2 | **0.2462** |
| ENSMUSG00000031878 | Nae1 | ENSMUSG00000000902 | Smarcb1 | **0.2444** |
| ENSMUSG00000029176 | Anapc4 | ENSMUSG00000000001 | Gnai3 | **0.242** |
| ENSMUSG00000035683 | Melk | ENSMUSG00000002297 | Dbf4 | **0.2385** |
| ENSMUSG00000034023 | Fancd2 | ENSMUSG00000024293 | Esco1 | **0.2383** |
| ENSMUSG00000039994 | Timeless | ENSMUSG00000018509 | Cenpv | **0.2367** |
| ENSMUSG00000035683 | Melk | ENSMUSG00000035439 | Haus8 | **0.2334** |
| ENSMUSG00000066149 | Cdc26 | ENSMUSG00000002068 | Ccne1 | **0.2319** |
| ENSMUSG00000039130 | Zc3hc1 | ENSMUSG00000032218 | Ccnb2 | **0.2271** |
| ENSMUSG00000040760 | Appl1 | ENSMUSG00000034021 | Pds5b | **0.2267** |
| ENSMUSG00000034906 | Ncaph | ENSMUSG00000034023 | Fancd2 | **0.226** |
| ENSMUSG00000069631 | Strada | ENSMUSG00000033952 | Aspm | **0.2236** |
| ENSMUSG00000026491 | Ahctf1 | ENSMUSG00000000708 | Kat2b | **0.2193** |
| ENSMUSG00000029910 | Mad2l1 | ENSMUSG00000010342 | Tex14 | **0.2184** |
| ENSMUSG00000040250 | Spata30 | ENSMUSG00000032113 | Chek1 | **0.2184** |
| ENSMUSG00000032113 | Chek1 | ENSMUSG00000000028 | Cdc45 | **0.2182** |
| ENSMUSG00000032477 | Cdc25a | ENSMUSG00000025907 | Rb1cc1 | **0.2167** |
| ENSMUSG00000038482 | Tfdp1 | ENSMUSG00000038416 | Cdc16 | **0.2141** |
| ENSMUSG00000036672 | Cenpt | ENSMUSG00000018509 | Cenpv | **0.2135** |
| ENSMUSG00000032218 | Ccnb2 | ENSMUSG00000029472 | Anapc5 | **0.2126** |
| ENSMUSG00000072980 | Oip5 | ENSMUSG00000029176 | Anapc4 | **0.2107** |
| ENSMUSG00000036928 | Stag3 | ENSMUSG00000024293 | Esco1 | **0.2098** |
| ENSMUSG00000038619 | Ensa | ENSMUSG00000035683 | Melk | **0.2088** |
| ENSMUSG00000039187 | Fanci | ENSMUSG00000031660 | Brd7 | **0.2074** |
| ENSMUSG00000046591 | Ticrr | ENSMUSG00000029472 | Anapc5 | **0.2067** |
| ENSMUSG00000069910 | Ccdc99 | ENSMUSG00000035024 | Ncapd3 | **0.206** |
| ENSMUSG00000029253 | Cenpc1 | ENSMUSG00000011960 | Ccnt1 | **0.2057** |
| ENSMUSG00000072980 | Oip5 | ENSMUSG00000029253 | Cenpc1 | **0.2042** |
| ENSMUSG00000033417 | Cacul1 | ENSMUSG00000002870 | Mcm2 | **0.2038** |
| ENSMUSG00000039130 | Zc3hc1 | ENSMUSG00000034165 | Ccnd3 | **0.2038** |
| ENSMUSG00000038619 | Ensa | ENSMUSG00000019917 | Sep-10 | **0.2036** |
| ENSMUSG00000047534 | Mis18bp1 | ENSMUSG00000028059 | Arhgef2 | **0.2032** |
| ENSMUSG00000025480 | Syce1 | ENSMUSG00000000743 | Chmp1a | **0.2027** |
| ENSMUSG00000038482 | Tfdp1 | ENSMUSG00000034165 | Ccnd3 | **0.2007** |
| ENSMUSG00000029283 | Cdc7 | ENSMUSG00000022141 | Nipbl | **0.1981** |
| ENSMUSG00000079562 | Maea | ENSMUSG00000027306 | Nusap1 | **0.1981** |
| ENSMUSG00000035437 | Rabgap1 | ENSMUSG00000029062 | Cdk11b | **0.196** |
| ENSMUSG00000069910 | Ccdc99 | ENSMUSG00000021635 | Rad17 | **0.1941** |
| ENSMUSG00000028896 | Rcc1 | ENSMUSG00000018821 | Avpi1 | **0.1931** |
| ENSMUSG00000061607 | Mdc1 | ENSMUSG00000023015 | Racgap1 | **0.1923** |
| ENSMUSG00000039994 | Timeless | ENSMUSG00000019942 | Cdk1 | **0.1917** |
| ENSMUSG00000038482 | Tfdp1 | ENSMUSG00000022070 | Bora | **0.1916** |
| ENSMUSG00000032264 | Zw10 | ENSMUSG00000021595 | Nsun2 | **0.1914** |
| ENSMUSG00000064302 | Clasp1 | ENSMUSG00000035683 | Melk | **0.1903** |
| ENSMUSG00000069631 | Strada | ENSMUSG00000039187 | Fanci | **0.1883** |
| ENSMUSG00000037286 | Stag1 | ENSMUSG00000034154 | Ino80 | **0.1877** |
| ENSMUSG00000028678 | Kif2c | ENSMUSG00000000028 | Cdc45 | **0.1875** |
| ENSMUSG00000021375 | Kif13a | ENSMUSG00000020107 | Anapc16 | **0.1869** |
| ENSMUSG00000030649 | 3200002M19Rik | ENSMUSG00000025862 | Stag2 | **0.1865** |
| ENSMUSG00000062234 | Gak | ENSMUSG00000031660 | Brd7 | **0.1865** |
| ENSMUSG00000079555 | Haus3 | ENSMUSG00000061607 | Mdc1 | **0.1858** |
| ENSMUSG00000030105 | Arl8b | ENSMUSG00000004591 | Pkn2 | **0.1846** |
| ENSMUSG00000040034 | Nup43 | ENSMUSG00000005410 | Mcm5 | **0.1836** |
| ENSMUSG00000072082 | Ccnf | ENSMUSG00000024056 | Ndc80 | **0.1835** |
| ENSMUSG00000036777 | Anln | ENSMUSG00000021548 | Ccnh | **0.1834** |
| ENSMUSG00000073705 | Apitd1 | ENSMUSG00000066149 | Cdc26 | **0.1831** |
| ENSMUSG00000029004 | Mll5 | ENSMUSG00000017146 | Brca1 | **0.1827** |
| ENSMUSG00000029176 | Anapc4 | ENSMUSG00000006398 | Cdc20 | **0.1824** |
| ENSMUSG00000048668 | Rhno1 | ENSMUSG00000018509 | Cenpv | **0.1818** |
| ENSMUSG00000046591 | Ticrr | ENSMUSG00000029283 | Cdc7 | **0.1814** |
| ENSMUSG00000012443 | Kif11 | ENSMUSG00000002068 | Ccne1 | **0.1803** |
| ENSMUSG00000023067 | Cdkn1a | ENSMUSG00000018509 | Cenpv | **0.1801** |
| ENSMUSG00000017499 | Cdc6 | ENSMUSG00000002068 | Ccne1 | **0.1795** |
| ENSMUSG00000006398 | Cdc20 | ENSMUSG00000002297 | Dbf4 | **0.1792** |
| ENSMUSG00000037544 | Dlgap5 | ENSMUSG00000025925 | Terf1 | **0.1785** |
| ENSMUSG00000040084 | Bub1b | ENSMUSG00000014859 | E2f4 | **0.1782** |
| ENSMUSG00000068744 | Psrc1 | ENSMUSG00000038393 | Txnip | **0.1779** |
| ENSMUSG00000036672 | Cenpt | ENSMUSG00000000743 | Chmp1a | **0.1775** |
| ENSMUSG00000032218 | Ccnb2 | ENSMUSG00000031878 | Nae1 | **0.1769** |
| ENSMUSG00000027550 | Lrrcc1 | ENSMUSG00000023940 | Sgol1 | **0.1766** |
| ENSMUSG00000045328 | Cenpe | ENSMUSG00000020107 | Anapc16 | **0.1764** |
| ENSMUSG00000029147 | Ppm1g | ENSMUSG00000002068 | Ccne1 | **0.1762** |
| ENSMUSG00000028678 | Kif2c | ENSMUSG00000025862 | Stag2 | **0.1761** |
| ENSMUSG00000059586 | Nsmce2 | ENSMUSG00000020415 | Pttg1 | **0.1761** |
| ENSMUSG00000061665 | Cd2ap | ENSMUSG00000029253 | Cenpc1 | **0.1751** |
| ENSMUSG00000004591 | Pkn2 | ENSMUSG00000000708 | Kat2b | **0.175** |
| ENSMUSG00000027379 | Bub1 | ENSMUSG00000020307 | Cdc34 | **0.1748** |
| ENSMUSG00000078652 | Psme3 | ENSMUSG00000000001 | Gnai3 | **0.1747** |
| ENSMUSG00000022070 | Bora | ENSMUSG00000018509 | Cenpv | **0.1733** |
| ENSMUSG00000034165 | Ccnd3 | ENSMUSG00000026779 | Mastl | **0.1732** |
| ENSMUSG00000029414 | Kntc1 | ENSMUSG00000019794 | Katna1 | **0.1731** |
| ENSMUSG00000038416 | Cdc16 | ENSMUSG00000002870 | Mcm2 | **0.1726** |
| ENSMUSG00000030641 | 4632434I11Rik | ENSMUSG00000021103 | Mnat1 | **0.1724** |
| ENSMUSG00000029910 | Mad2l1 | ENSMUSG00000023505 | Cdca3 | **0.1723** |
| ENSMUSG00000047534 | Mis18bp1 | ENSMUSG00000033952 | Aspm | **0.1723** |
| ENSMUSG00000061607 | Mdc1 | ENSMUSG00000033952 | Aspm | **0.1721** |
| ENSMUSG00000029003 | Mad2l2 | ENSMUSG00000026361 | Cdc73 | **-0.1738** |
| ENSMUSG00000042029 | Ncapg2 | ENSMUSG00000038393 | Txnip | **-0.1767** |
| ENSMUSG00000072980 | Oip5 | ENSMUSG00000033054 | Npat | **-0.1851** |
| ENSMUSG00000017499 | Cdc6 | ENSMUSG00000000743 | Chmp1a | **-0.1859** |
| ENSMUSG00000004085 | B230120H23Rik | ENSMUSG00000002068 | Ccne1 | **-0.1921** |
| ENSMUSG00000022945 | Chaf1b | ENSMUSG00000019794 | Katna1 | **-0.1941** |
| ENSMUSG00000038482 | Tfdp1 | ENSMUSG00000004085 | B230120H23Rik | **-0.2037** |
| ENSMUSG00000033952 | Aspm | ENSMUSG00000000743 | Chmp1a | **-0.2117** |
| ENSMUSG00000042688 | Mapk6 | ENSMUSG00000022945 | Chaf1b | **-0.2119** |
| ENSMUSG00000066357 | Wdr6 | ENSMUSG00000021635 | Rad17 | **-0.22** |
| ENSMUSG00000032411 | Tfdp2 | ENSMUSG00000017499 | Cdc6 | **-0.2289** |
| ENSMUSG00000033952 | Aspm | ENSMUSG00000025480 | Syce1 | **-0.2346** |

Table S4: Top gene pairs in G1 stage only.

| ID 1 | Assoc. gene 1 | ID 2 | Assoc. gene 2 | $\rho_{G2M}$ |
|---|---|---|---|---|
| ENSMUSG00000036928 | Stag3 | ENSMUSG00000021276 | Cinp | **0.4194** |
| ENSMUSG00000038619 | Ensa | ENSMUSG00000029176 | Anapc4 | **0.3732** |
| ENSMUSG00000073705 | Apitd1 | ENSMUSG00000036928 | Stag3 | **0.3711** |
| ENSMUSG00000031371 | Haus7 | ENSMUSG00000026965 | Anapc2 | **0.3333** |
| ENSMUSG00000026779 | Mastl | ENSMUSG00000021965 | Ska3 | **0.3146** |
| ENSMUSG00000036928 | Stag3 | ENSMUSG00000031371 | Haus7 | **0.3023** |
| ENSMUSG00000023572 | Ccndbp1 | ENSMUSG00000018821 | Avpi1 | **0.2975** |
| ENSMUSG00000035828 | Pim3 | ENSMUSG00000031095 | Cul4b | **0.2836** |
| ENSMUSG00000034021 | Pds5b | ENSMUSG00000018509 | Cenpv | **0.2805** |
| ENSMUSG00000036782 | Klhl13 | ENSMUSG00000000743 | Chmp1a | **0.2746** |
| ENSMUSG00000032562 | Gnai2 | ENSMUSG00000027715 | Ccna2 | **0.274** |
| ENSMUSG00000028066 | Pmf1 | ENSMUSG00000026753 | Ppp6c | **0.2737** |
| ENSMUSG00000039187 | Fanci | ENSMUSG00000026965 | Anapc2 | **0.2737** |
| ENSMUSG00000038619 | Ensa | ENSMUSG00000036782 | Klhl13 | **0.2715** |
| ENSMUSG00000036928 | Stag3 | ENSMUSG00000035828 | Pim3 | **0.2664** |
| ENSMUSG00000036850 | Mrpl41 | ENSMUSG00000020415 | Pttg1 | **0.2604** |
| ENSMUSG00000036782 | Klhl13 | ENSMUSG00000023572 | Ccndbp1 | **0.2582** |
| ENSMUSG00000033054 | Npat | ENSMUSG00000021965 | Ska3 | **0.2556** |
| ENSMUSG00000039187 | Fanci | ENSMUSG00000025480 | Syce1 | **0.255** |
| ENSMUSG00000038619 | Ensa | ENSMUSG00000036850 | Mrpl41 | **0.251** |
| ENSMUSG00000028896 | Rcc1 | ENSMUSG00000021276 | Cinp | **0.2484** |
| ENSMUSG00000038416 | Cdc16 | ENSMUSG00000018509 | Cenpv | **0.248** |
| ENSMUSG00000035683 | Melk | ENSMUSG00000029910 | Mad2l1 | **0.2466** |
| ENSMUSG00000019923 | Zwint | ENSMUSG00000018821 | Avpi1 | **0.246** |
| ENSMUSG00000038416 | Cdc16 | ENSMUSG00000034023 | Fancd2 | **0.244** |
| ENSMUSG00000034023 | Fancd2 | ENSMUSG00000003824 | Syce2 | **0.2419** |
| ENSMUSG00000072980 | Oip5 | ENSMUSG00000029003 | Mad2l2 | **0.2415** |
| ENSMUSG00000020415 | Pttg1 | ENSMUSG00000006398 | Cdc20 | **0.2398** |
| ENSMUSG00000029472 | Anapc5 | ENSMUSG00000020107 | Anapc16 | **0.2396** |
| ENSMUSG00000031723 | Txnl4b | ENSMUSG00000022978 | Mis18a | **0.2391** |
| ENSMUSG00000021276 | Cinp | ENSMUSG00000020415 | Pttg1 | **0.2386** |
| ENSMUSG00000073705 | Apitd1 | ENSMUSG00000041997 | Tlk1 | **0.2384** |
| ENSMUSG00000036928 | Stag3 | ENSMUSG00000028896 | Rcc1 | **0.2382** |
| ENSMUSG00000042029 | Ncapg2 | ENSMUSG00000027363 | Usp8 | **0.2382** |
| ENSMUSG00000053119 | Chmp3 | ENSMUSG00000022978 | Mis18a | **0.2377** |
| ENSMUSG00000029283 | Cdc7 | ENSMUSG00000025616 | Usp16 | **0.2356** |
| ENSMUSG00000031371 | Haus7 | ENSMUSG00000000743 | Chmp1a | **0.2323** |
| ENSMUSG00000039187 | Fanci | ENSMUSG00000028896 | Rcc1 | **0.2316** |
| ENSMUSG00000039187 | Fanci | ENSMUSG00000025616 | Usp16 | **0.2313** |
| ENSMUSG00000034165 | Ccnd3 | ENSMUSG00000000743 | Chmp1a | **0.2302** |
| ENSMUSG00000036928 | Stag3 | ENSMUSG00000000902 | Smarcb1 | **0.2302** |
| ENSMUSG00000026779 | Mastl | ENSMUSG00000025480 | Syce1 | **0.2272** |
| ENSMUSG00000029003 | Mad2l2 | ENSMUSG00000019794 | Katna1 | **0.2261** |
| ENSMUSG00000036672 | Cenpt | ENSMUSG00000019923 | Zwint | **0.2261** |
| ENSMUSG00000036928 | Stag3 | ENSMUSG00000036779 | Papd5 | **0.2258** |
| ENSMUSG00000036928 | Stag3 | ENSMUSG00000023572 | Ccndbp1 | **0.2254** |
| ENSMUSG00000079555 | Haus3 | ENSMUSG00000000708 | Kat2b | **0.2244** |
| ENSMUSG00000036782 | Klhl13 | ENSMUSG00000021276 | Cinp | **0.2237** |
| ENSMUSG00000034165 | Ccnd3 | ENSMUSG00000032264 | Zw10 | **0.2226** |
| ENSMUSG00000039187 | Fanci | ENSMUSG00000023572 | Ccndbp1 | **0.2222** |
| ENSMUSG00000040034 | Nup43 | ENSMUSG00000029003 | Mad2l2 | **0.2214** |
| ENSMUSG00000029176 | Anapc4 | ENSMUSG00000018821 | Avpi1 | **0.2212** |
| ENSMUSG00000068744 | Psrc1 | ENSMUSG00000037725 | Ckap2 | **0.2193** |
| ENSMUSG00000041133 | Smc1a | ENSMUSG00000028678 | Kif2c | **0.2185** |
| ENSMUSG00000028447 | Dctn3 | ENSMUSG00000022945 | Chaf1b | **0.2183** |
| ENSMUSG00000036850 | Mrpl41 | ENSMUSG00000036782 | Klhl13 | **0.2166** |
| ENSMUSG00000036779 | Papd5 | ENSMUSG00000019923 | Zwint | **0.2164** |
| ENSMUSG00000038619 | Ensa | ENSMUSG00000018509 | Cenpv | **0.2158** |
| ENSMUSG00000074476 | Spc24 | ENSMUSG00000019923 | Zwint | **0.2158** |
| ENSMUSG00000035437 | Rabgap1 | ENSMUSG00000022678 | Nde1 | **0.2156** |
| ENSMUSG00000038393 | Txnip | ENSMUSG00000023572 | Ccndbp1 | **0.215** |
| ENSMUSG00000029003 | Mad2l2 | ENSMUSG00000020516 | Rps6kb1 | **0.2133** |
| ENSMUSG00000023067 | Cdkn1a | ENSMUSG00000018821 | Avpi1 | **0.2129** |
| ENSMUSG00000066149 | Cdc26 | ENSMUSG00000023940 | Sgol1 | **0.2126** |
| ENSMUSG00000036782 | Klhl13 | ENSMUSG00000026753 | Ppp6c | **0.2108** |
| ENSMUSG00000020415 | Pttg1 | ENSMUSG00000019923 | Zwint | **0.2101** |
| ENSMUSG00000036782 | Klhl13 | ENSMUSG00000021965 | Ska3 | **0.2097** |
| ENSMUSG00000034165 | Ccnd3 | ENSMUSG00000031371 | Haus7 | **0.2096** |
| ENSMUSG00000025480 | Syce1 | ENSMUSG00000000743 | Chmp1a | **0.2094** |
| ENSMUSG00000025480 | Syce1 | ENSMUSG00000021965 | Ska3 | **0.2092** |
| ENSMUSG00000027285 | Haus2 | ENSMUSG00000021548 | Ccnh | **0.2092** |
| ENSMUSG00000038619 | Ensa | ENSMUSG00000026491 | Ahctf1 | **0.2077** |
| ENSMUSG00000073705 | Apitd1 | ENSMUSG00000038393 | Txnip | **0.2067** |
| ENSMUSG00000036782 | Klhl13 | ENSMUSG00000029176 | Anapc4 | **0.2053** |
| ENSMUSG00000034906 | Ncaph | ENSMUSG00000023572 | Ccndbp1 | **0.2046** |
| ENSMUSG00000028896 | Rcc1 | ENSMUSG00000020415 | Pttg1 | **0.2038** |
| ENSMUSG00000036850 | Mrpl41 | ENSMUSG00000027285 | Haus2 | **0.2032** |
| ENSMUSG00000062234 | Gak | ENSMUSG00000036782 | Klhl13 | **0.2031** |
| ENSMUSG00000021276 | Cinp | ENSMUSG00000000708 | Kat2b | **0.203** |
| ENSMUSG00000020415 | Pttg1 | ENSMUSG00000000708 | Kat2b | **0.2027** |
| ENSMUSG00000046591 | Ticrr | ENSMUSG00000021635 | Rad17 | **0.2024** |
| ENSMUSG00000073705 | Apitd1 | ENSMUSG00000031371 | Haus7 | **0.2018** |
| ENSMUSG00000036850 | Mrpl41 | ENSMUSG00000021548 | Ccnh | **0.2017** |
| ENSMUSG00000062234 | Gak | ENSMUSG00000021965 | Ska3 | **0.2017** |
| ENSMUSG00000032264 | Zw10 | ENSMUSG00000026361 | Cdc73 | **0.2016** |
| ENSMUSG00000034165 | Ccnd3 | ENSMUSG00000021965 | Ska3 | **0.2015** |
| ENSMUSG00000031371 | Haus7 | ENSMUSG00000021965 | Ska3 | **0.2011** |
| ENSMUSG00000036850 | Mrpl41 | ENSMUSG00000034021 | Pds5b | **0.201** |
| ENSMUSG00000038416 | Cdc16 | ENSMUSG00000029521 | Chek2 | **0.2003** |
| ENSMUSG00000029176 | Anapc4 | ENSMUSG00000028896 | Rcc1 | **0.1996** |
| ENSMUSG00000031723 | Txnl4b | ENSMUSG00000020415 | Pttg1 | **0.1988** |
| ENSMUSG00000039130 | Zc3hc1 | ENSMUSG00000036672 | Cenpt | **0.1977** |
| ENSMUSG00000042750 | Bex2 | ENSMUSG00000039130 | Zc3hc1 | **0.1977** |
| ENSMUSG00000030105 | Arl8b | ENSMUSG00000027469 | Tpx2 | **0.1976** |
| ENSMUSG00000039130 | Zc3hc1 | ENSMUSG00000036782 | Klhl13 | **0.1974** |
| ENSMUSG00000036782 | Klhl13 | ENSMUSG00000024989 | Cep55 | **-0.2022** |
| ENSMUSG00000036782 | Klhl13 | ENSMUSG00000025616 | Usp16 | **-0.2221** |
| ENSMUSG00000061607 | Mdc1 | ENSMUSG00000026779 | Mastl | **-0.228** |
| ENSMUSG00000079555 | Haus3 | ENSMUSG00000038416 | Cdc16 | **-0.2355** |
| ENSMUSG00000039187 | Fanci | ENSMUSG00000033417 | Cacul1 | **-0.2453** |

Table S5: Top gene pairs in G2M stage only.

| ID 1 | Assoc. gene 1 | ID 2 | Assoc. gene 2 | $\rho_S$ |
|---|---|---|---|---|
| ENSMUSG00000028059 | Arhgef2 | ENSMUSG00000025862 | Stag2 | **0.6874** |
| ENSMUSG00000034218 | Atm | ENSMUSG00000028059 | Arhgef2 | **0.6812** |
| ENSMUSG00000032113 | Chek1 | ENSMUSG00000028059 | Arhgef2 | **0.6667** |
| ENSMUSG00000033054 | Npat | ENSMUSG00000032113 | Chek1 | **0.6316** |
| ENSMUSG00000032113 | Chek1 | ENSMUSG00000031858 | Mau2 | **0.6293** |
| ENSMUSG00000029176 | Anapc4 | ENSMUSG00000028059 | Arhgef2 | **0.6268** |
| ENSMUSG00000034218 | Atm | ENSMUSG00000026779 | Mastl | **0.6005** |
| ENSMUSG00000034218 | Atm | ENSMUSG00000032113 | Chek1 | **0.5869** |
| ENSMUSG00000033054 | Npat | ENSMUSG00000028059 | Arhgef2 | **0.5868** |
| ENSMUSG00000029176 | Anapc4 | ENSMUSG00000025862 | Stag2 | **0.5803** |
| ENSMUSG00000031858 | Mau2 | ENSMUSG00000028059 | Arhgef2 | **0.5783** |
| ENSMUSG00000066149 | Cdc26 | ENSMUSG00000000708 | Kat2b | **0.5754** |
| ENSMUSG00000032113 | Chek1 | ENSMUSG00000029501 | Ankle2 | **0.5749** |
| ENSMUSG00000066149 | Cdc26 | ENSMUSG00000029253 | Cenpc1 | **0.5738** |
| ENSMUSG00000032113 | Chek1 | ENSMUSG00000025862 | Stag2 | **0.5679** |
| ENSMUSG00000034218 | Atm | ENSMUSG00000010342 | Tex14 | **0.5573** |
| ENSMUSG00000045969 | Ing1 | ENSMUSG00000028059 | Arhgef2 | **0.5558** |
| ENSMUSG00000057110 | Cep110 | ENSMUSG00000000001 | Gnai3 | **0.5515** |
| ENSMUSG00000033054 | Npat | ENSMUSG00000031858 | Mau2 | **0.5476** |
| ENSMUSG00000038393 | Txnip | ENSMUSG00000032113 | Chek1 | **0.5449** |
| ENSMUSG00000031858 | Mau2 | ENSMUSG00000025862 | Stag2 | **0.5384** |
| ENSMUSG00000032113 | Chek1 | ENSMUSG00000029176 | Anapc4 | **0.5354** |
| ENSMUSG00000041238 | Rbbp8 | ENSMUSG00000029501 | Ankle2 | **0.5325** |
| ENSMUSG00000029176 | Anapc4 | ENSMUSG00000029147 | Ppm1g | **0.5288** |
| ENSMUSG00000030649 | 3200002M19Rik | ENSMUSG00000010342 | Tex14 | **0.5284** |
| ENSMUSG00000031371 | Haus7 | ENSMUSG00000029501 | Ankle2 | **0.5222** |
| ENSMUSG00000032113 | Chek1 | ENSMUSG00000022945 | Chaf1b | **0.5203** |
| ENSMUSG00000057110 | Cep110 | ENSMUSG00000026779 | Mastl | **0.5198** |
| ENSMUSG00000032113 | Chek1 | ENSMUSG00000030641 | 4632434I11Rik | **0.517** |
| ENSMUSG00000057110 | Cep110 | ENSMUSG00000010342 | Tex14 | **0.5168** |
| ENSMUSG00000066149 | Cdc26 | ENSMUSG00000020107 | Anapc16 | **0.5152** |
| ENSMUSG00000034218 | Atm | ENSMUSG00000030649 | 3200002M19Rik | **0.514** |
| ENSMUSG00000031858 | Mau2 | ENSMUSG00000029176 | Anapc4 | **0.5129** |
| ENSMUSG00000038393 | Txnip | ENSMUSG00000029501 | Ankle2 | **0.5114** |
| ENSMUSG00000045328 | Cenpe | ENSMUSG00000028678 | Kif2c | **0.5106** |
| ENSMUSG00000030641 | 4632434I11Rik | ENSMUSG00000029003 | Mad2l2 | **0.5089** |
| ENSMUSG00000066149 | Cdc26 | ENSMUSG00000020694 | Tlk2 | **0.5087** |
| ENSMUSG00000029253 | Cenpc1 | ENSMUSG00000000708 | Kat2b | **0.5073** |
| ENSMUSG00000038393 | Txnip | ENSMUSG00000030061 | Uba3 | **0.5072** |
| ENSMUSG00000045969 | Ing1 | ENSMUSG00000032113 | Chek1 | **0.506** |
| ENSMUSG00000034218 | Atm | ENSMUSG00000029176 | Anapc4 | **0.5059** |
| ENSMUSG00000057110 | Cep110 | ENSMUSG00000034218 | Atm | **0.5042** |
| ENSMUSG00000029501 | Ankle2 | ENSMUSG00000014355 | Anapc1 | **0.501** |
| ENSMUSG00000039128 | Cdc123 | ENSMUSG00000023572 | Ccndbp1 | **0.5008** |
| ENSMUSG00000026779 | Mastl | ENSMUSG00000010342 | Tex14 | **0.5003** |
| ENSMUSG00000033054 | Npat | ENSMUSG00000029176 | Anapc4 | **0.498** |
| ENSMUSG00000034218 | Atm | ENSMUSG00000031371 | Haus7 | **0.4969** |
| ENSMUSG00000045969 | Ing1 | ENSMUSG00000029176 | Anapc4 | **0.495** |
| ENSMUSG00000057110 | Cep110 | ENSMUSG00000014859 | E2f4 | **0.4946** |
| ENSMUSG00000031858 | Mau2 | ENSMUSG00000029147 | Ppm1g | **0.4934** |
| ENSMUSG00000034218 | Atm | ENSMUSG00000022945 | Chaf1b | **0.4934** |
| ENSMUSG00000029253 | Cenpc1 | ENSMUSG00000020694 | Tlk2 | **0.492** |
| ENSMUSG00000032113 | Chek1 | ENSMUSG00000023572 | Ccndbp1 | **0.4908** |
| ENSMUSG00000034218 | Atm | ENSMUSG00000025862 | Stag2 | **0.4904** |
| ENSMUSG00000032113 | Chek1 | ENSMUSG00000014355 | Anapc1 | **0.4897** |
| ENSMUSG00000069631 | Strada | ENSMUSG00000045969 | Ing1 | **0.489** |
| ENSMUSG00000032113 | Chek1 | ENSMUSG00000029003 | Mad2l2 | **0.4883** |
| ENSMUSG00000014859 | E2f4 | ENSMUSG00000000708 | Kat2b | **0.4876** |
| ENSMUSG00000038619 | Ensa | ENSMUSG00000032113 | Chek1 | **0.484** |
| ENSMUSG00000045969 | Ing1 | ENSMUSG00000025862 | Stag2 | **0.481** |
| ENSMUSG00000030641 | 4632434I11Rik | ENSMUSG00000021595 | Nsun2 | **0.48** |
| ENSMUSG00000029501 | Ankle2 | ENSMUSG00000029003 | Mad2l2 | **0.4787** |
| ENSMUSG00000029147 | Ppm1g | ENSMUSG00000028059 | Arhgef2 | **0.4782** |
| ENSMUSG00000029501 | Ankle2 | ENSMUSG00000022945 | Chaf1b | **0.4761** |
| ENSMUSG00000010342 | Tex14 | ENSMUSG00000000001 | Gnai3 | **0.4757** |
| ENSMUSG00000029147 | Ppm1g | ENSMUSG00000025862 | Stag2 | **0.475** |
| ENSMUSG00000030649 | 3200002M19Rik | ENSMUSG00000026779 | Mastl | **0.4733** |
| ENSMUSG00000014859 | E2f4 | ENSMUSG00000000001 | Gnai3 | **0.4732** |
| ENSMUSG00000034218 | Atm | ENSMUSG00000029501 | Ankle2 | **0.4715** |
| ENSMUSG00000030061 | Uba3 | ENSMUSG00000029501 | Ankle2 | **0.4706** |
| ENSMUSG00000038393 | Txnip | ENSMUSG00000020107 | Anapc16 | **0.4681** |
| ENSMUSG00000026779 | Mastl | ENSMUSG00000000001 | Gnai3 | **0.4653** |
| ENSMUSG00000066149 | Cdc26 | ENSMUSG00000038252 | Ncapd2 | **0.46** |
| ENSMUSG00000033054 | Npat | ENSMUSG00000029501 | Ankle2 | **0.4597** |
| ENSMUSG00000029501 | Ankle2 | ENSMUSG00000028873 | Cdca8 | **0.4596** |
| ENSMUSG00000038252 | Ncapd2 | ENSMUSG00000020694 | Tlk2 | **0.4592** |
| ENSMUSG00000045969 | Ing1 | ENSMUSG00000031858 | Mau2 | **0.4592** |
| ENSMUSG00000029003 | Mad2l2 | ENSMUSG00000023572 | Ccndbp1 | **0.4579** |
| ENSMUSG00000045328 | Cenpe | ENSMUSG00000038393 | Txnip | **0.4552** |
| ENSMUSG00000032113 | Chek1 | ENSMUSG00000021548 | Ccnh | **0.4547** |
| ENSMUSG00000041238 | Rbbp8 | ENSMUSG00000038393 | Txnip | **0.4539** |
| ENSMUSG00000030641 | 4632434I11Rik | ENSMUSG00000023572 | Ccndbp1 | **0.4537** |
| ENSMUSG00000033054 | Npat | ENSMUSG00000029147 | Ppm1g | **0.4537** |
| ENSMUSG00000061665 | Cd2ap | ENSMUSG00000038393 | Txnip | **0.4531** |
| ENSMUSG00000034218 | Atm | ENSMUSG00000029003 | Mad2l2 | **0.4523** |
| ENSMUSG00000041238 | Rbbp8 | ENSMUSG00000030641 | 4632434I11Rik | **0.4517** |
| ENSMUSG00000034218 | Atm | ENSMUSG00000033054 | Npat | **0.4514** |
| ENSMUSG00000033417 | Cacul1 | ENSMUSG00000000001 | Gnai3 | **0.4501** |
| ENSMUSG00000038393 | Txnip | ENSMUSG00000031858 | Mau2 | **0.4499** |
| ENSMUSG00000040084 | Bub1b | ENSMUSG00000021103 | Mnat1 | **0.4497** |
| ENSMUSG00000042750 | Bex2 | ENSMUSG00000034218 | Atm | **0.4495** |
| ENSMUSG00000036782 | Klhl13 | ENSMUSG00000034218 | Atm | **0.4484** |
| ENSMUSG00000034218 | Atm | ENSMUSG00000030641 | 4632434I11Rik | **0.448** |
| ENSMUSG00000078652 | Psme3 | ENSMUSG00000045328 | Cenpe | **0.4475** |
| ENSMUSG00000039128 | Cdc123 | ENSMUSG00000029003 | Mad2l2 | **0.447** |
| ENSMUSG00000029501 | Ankle2 | ENSMUSG00000020107 | Anapc16 | **0.4465** |
| ENSMUSG00000041238 | Rbbp8 | ENSMUSG00000032113 | Chek1 | **0.4462** |
| ENSMUSG00000045328 | Cenpe | ENSMUSG00000017146 | Brca1 | **0.4462** |
| ENSMUSG00000057110 | Cep110 | ENSMUSG00000005410 | Mcm5 | **0.4447** |
| ENSMUSG00000032504 | Pdcd6ip | ENSMUSG00000002635 | Pdcd2l | **0.4444** |

Table S6: Top gene pairs in S stage only.

### 7 Single-cell RNA data analysis results

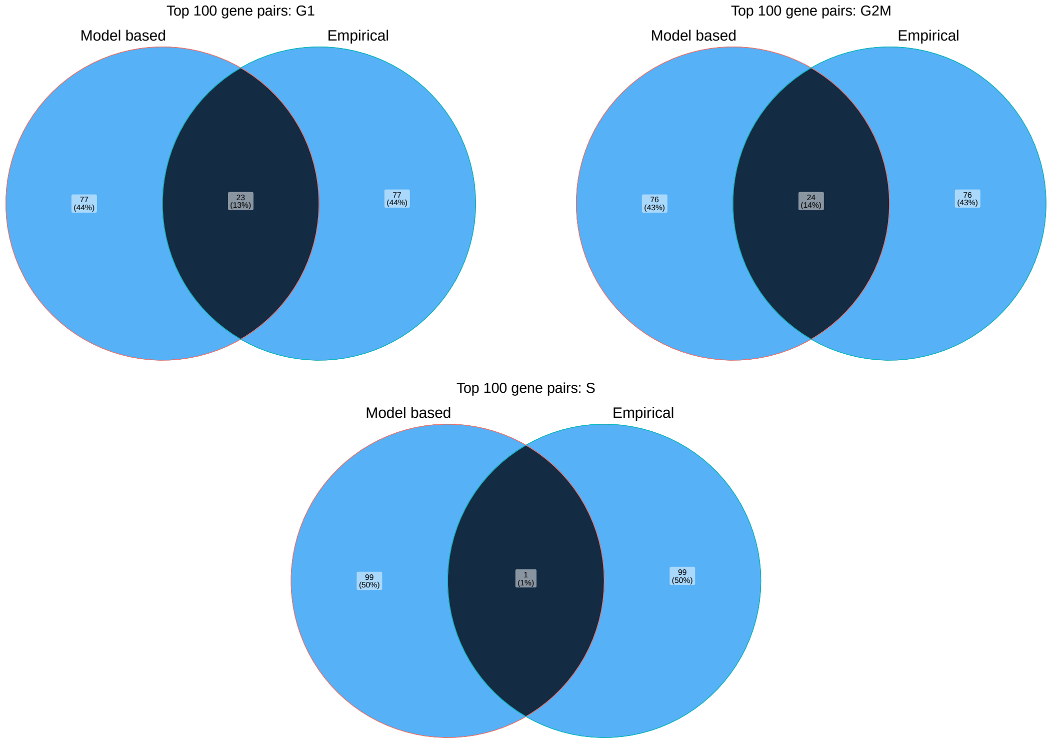


Figure S7: Overlap in top 100 $\rho$ values between model based and empirical estimates in G1, G2M, and S stages.

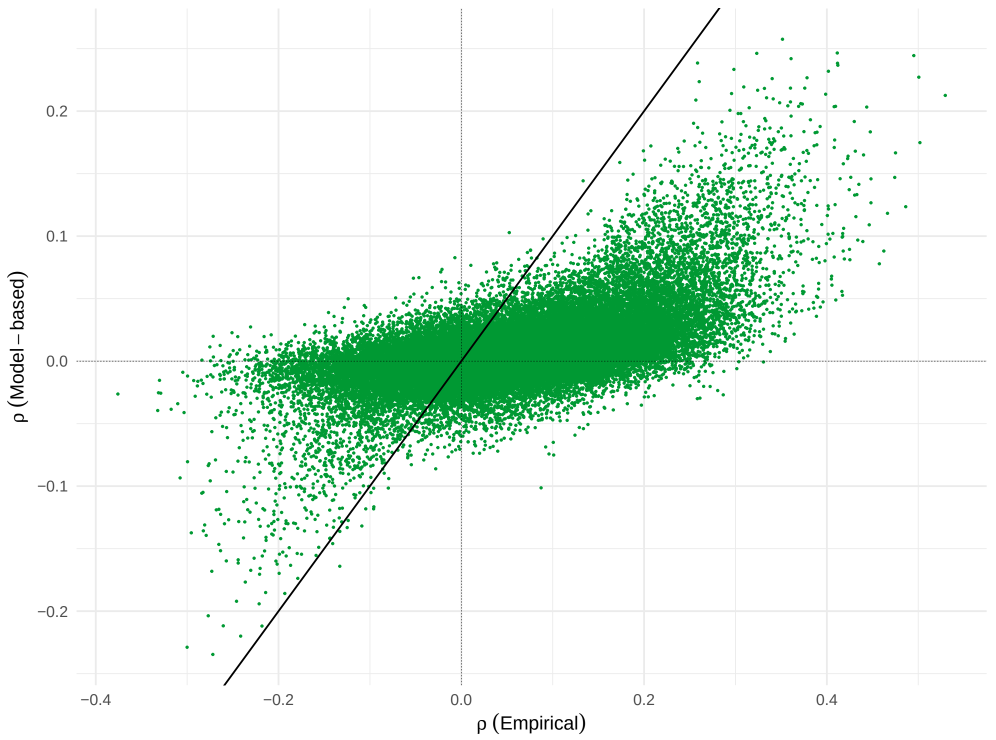


Figure S8: Comparing model-based and empirical estimates of $\rho$ in G1 phase in the mESC dataset.

### 8 Pediatric-onset MS metagenomic data analysis results

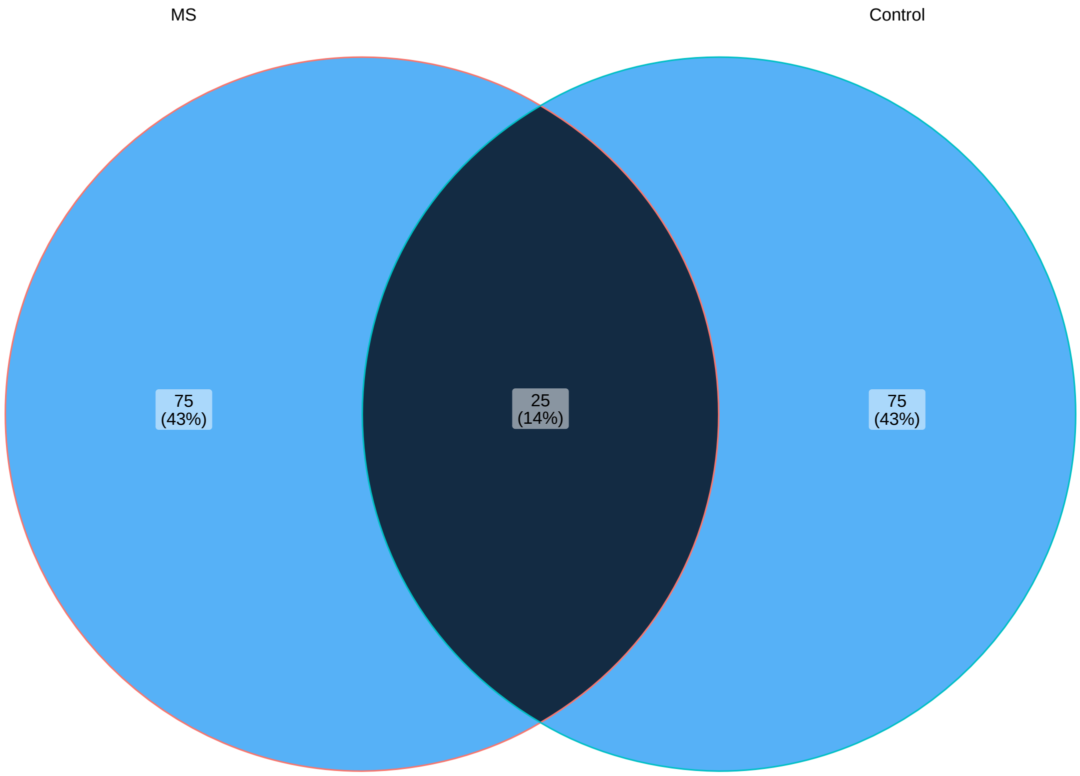




Figure S9: Overlap in top 100 $\rho$ values between model based and empirical estimates in MS vs. unaffected controls.

| Genus 1 | Genus 2 | $\rho_{MS}$ |
|---|---|---|
| Phil12 | BacteroidesA | 0.6385 |
| RuminiclostridiumE | RuminococcusE | 0.6226 |
| CAG-170 | CAG-180 | 0.6126 |
| BacteroidesB | BacteroidesA | 0.612 |
| Butyricimonas | BacteroidesB | 0.6093 |
| 43-108 | BacteroidesA | 0.6073 |
| OlsenellaE | Collinsella | 0.5942 |
| CAG-1427 | UBA11524 | 0.5895 |
| Senegalimassilia | OlsenellaE | 0.5838 |
| Phil12 | BacteroidesB | 0.5822 |
| Butyricimonas | Prevotella | 0.5779 |
| Paraprevotella | BacteroidesB | 0.5641 |
| Phil12 | 43-108 | 0.5618 |
| Gabonibacter | 43-108 | 0.5612 |
| Pauljensenia | Collinsella | 0.5581 |
| Turicibacter | Romboutsia | 0.5465 |
| AlistipesA | CAG-177 | 0.5418 |
| Adlercreutzia | EubacteriumR | 0.5414 |
| CAG-170 | CAG-83 | 0.5408 |
| 43-108 | BacteroidesB | 0.5344 |
| Christensenella | UBA1191 | 0.5334 |
| 43-108 | Butyricimonas | 0.5332 |
| AlistipesA | UBA11524 | 0.5317 |
| Senegalimassilia | Collinsella | 0.5317 |
| PeH17 | CAG-1427 | 0.5248 |
| UBA1191 | CAG-170 | 0.5243 |
| Intestinibacter | Romboutsia | 0.5211 |
| Gabonibacter | Butyricimonas | 0.5198 |
| UBA11774 | RuminiclostridiumC | 0.5192 |
| Gabonibacter | BacteroidesA | 0.5138 |
| Phil12 | Butyricimonas | 0.5112 |
| Butyricimonas | BacteroidesA | 0.5094 |
| RuminiclostridiumE | UBA1417 | 0.5041 |
| RuminiclostridiumC | Adlercreutzia | 0.5026 |
| Gabonibacter | Prevotella | 0.4961 |
| UBA11774 | RuminococcusA | 0.4959 |
| Coprobacillus | CAG-180 | 0.493 |
| PeH17 | CAG-177 | 0.4928 |
| Paraprevotella | Butyricimonas | 0.4924 |
| CAG-103 | CAG-177 | 0.4918 |
| CAG-103 | UBA11524 | 0.4915 |
| PeH17 | CAG-170 | 0.491 |
| Acutalibacter | UBA1417 | 0.4885 |
| Coprobacillus | PeH17 | 0.4855 |
| CAG-180 | CAG-83 | 0.4854 |
| Parabacteroides | BacteroidesA | 0.4851 |
| CAG-177 | CAG-83 | 0.4831 |
| Pauljensenia | OlsenellaE | 0.4815 |
| UBA11524 | RuminococcusD | 0.4811 |
| EubacteriumF | RuminococcusC | 0.4808 |
| CAG-103 | RuminococcusC | 0.4781 |
| Senegalimassilia | LactobacillusB | 0.4776 |
| Barnesiella | Dialister | 0.4772 |
| Parabacteroides | BacteroidesB | 0.4741 |
| UBA1191 | PeH17 | 0.474 |
| CAG-170 | CAG-110 | 0.4737 |
| Phil12 | Parabacteroides | 0.4728 |
| Christensenella | CAG-170 | 0.4724 |
| PeH17 | CAG-180 | 0.472 |
| UBA1191 | CAG-83 | 0.4706 |
| Gabonibacter | BacteroidesB | 0.4683 |
| PeH17 | CAG-103 | 0.4665 |
| Clostridium | RuminococcusE | 0.4624 |
| PeH17 | CAG-83 | 0.4624 |
| CAG-110 | CAG-83 | 0.4617 |
| Coprobacillus | CAG-170 | 0.4616 |
| Christensenella | CAG-177 | 0.4608 |
| Parabacteroides | Alistipes | 0.4597 |
| Butyricimonas | Parabacteroides | 0.4592 |
| PeH17 | Adlercreutzia | 0.4586 |
| Phil12 | Lachnospira | 0.4547 |
| Gabonibacter | Phil12 | 0.4546 |
| Christensenella | AlistipesA | 0.4542 |
| PeH17 | UBA11524 | 0.4526 |
| Escherichia | LactobacillusB | 0.4514 |
| Gabonibacter | Alistipes | 0.4502 |
| AlistipesA | RuminococcusD | 0.4497 |
| Gabonibacter | Parabacteroides | 0.4493 |
| Paraprevotella | BacteroidesA | 0.4486 |
| UBA1191 | CAG-1427 | 0.4474 |
| CAG-103 | Adlercreutzia | 0.445 |
| 43-108 | Parabacteroides | 0.4441 |
| Acutalibacter | ClostridiumA | 0.4427 |
| Holdemanella | LactobacillusB | 0.4424 |
| CAG-170 | CAG-177 | 0.4418 |
| RuminiclostridiumE | CAG-127 | 0.4397 |
| UBA1191 | CAG-177 | 0.4387 |
| Pauljensenia | Mogibacterium | 0.4385 |
| Barnesiella | RuminococcusD | 0.4384 |
| UBA1191 | UBA1417 | 0.4375 |
| Pauljensenia | Eggerthella | 0.4362 |
| Christensenella | Acutalibacter | 0.4335 |
| Phil12 | Paraprevotella | 0.4329 |
| Gordonibacter | Ruthenibacterium | 0.4317 |
| CAG-1427 | CAG-103 | 0.4305 |
| UBA1417 | Collinsella | 0.4302 |
| UBA11524 | CAG-177 | 0.43 |
| Pauljensenia | Acutalibacter | 0.4255 |
| Christensenella | CAG-180 | 0.4251 |
| Coprobacillus | UBA1191 | 0.4248 |

| Genus 1 | Genus 2 | $\rho_C$ |
|---|---|---|
| KLE1615 | Lachnospira | 0.6761 |
| Barnesiella | 43-108 | 0.6627 |
| Phil12 | BacteroidesA | 0.646 |
| CAG-103 | CAG-83 | 0.6413 |
| CAG-127 | RuminococcusC | 0.6368 |
| Acidaminococcus | CAG-127 | 0.6221 |
| Acidaminococcus | Megasphaera | 0.622 |
| CAG-177 | Holdemanella | 0.6056 |
| Intestinibacter | Clostridium | 0.6022 |
| RC9 | Phil12 | 0.5997 |
| CAG-170 | CAG-110 | 0.5948 |
| CAG-103 | UBA11524 | 0.5919 |
| Paraprevotella | Barnesiella | 0.59 |
| Paraprevotella | BacteroidesB | 0.5896 |
| UBA1191 | UBA1417 | 0.5891 |
| RC9 | Paraprevotella | 0.5881 |
| Acutalibacter | ClostridiumA | 0.5829 |
| RuminococcusC | RuminococcusD | 0.5807 |
| CAG-110 | CAG-83 | 0.5777 |
| PeH17 | CAG-170 | 0.5767 |
| PeH17 | Holdemanella | 0.5722 |
| UBA11774 | RuminococcusD | 0.569 |
| TF01-11 | Lachnospira | 0.5684 |
| CAG-127 | RuminococcusD | 0.5635 |
| UBA11774 | CAG-127 | 0.5632 |
| Acidaminococcus | RuminococcusC | 0.563 |
| OlsenellaE | Collinsella | 0.5627 |
| ER4 | CAG-83 | 0.5604 |
| Senegalimassilia | Bifidobacterium | 0.5569 |
| PeH17 | CAG-110 | 0.5562 |
| 43-108 | Alistipes | 0.5505 |
| CAG-180 | RuminococcusD | 0.5499 |
| RuminiclostridiumE | EubacteriumR | 0.5482 |
| RC9 | Gabonibacter | 0.5481 |
| TF01-11 | KLE1615 | 0.5471 |
| Sellimonas | CAG-81 | 0.5457 |
| RuminiclostridiumE | RuminococcusD | 0.5441 |
| Megasphaera | LactobacillusB | 0.5432 |
| Clostridium | Romboutsia | 0.5416 |
| RC9 | BacteroidesA | 0.5416 |
| RC9 | Oscillibacter | 0.5354 |
| RuminiclostridiumE | Megasphaera | 0.5342 |
| Phil12 | Butyricimonas | 0.5333 |
| Phil12 | Paraprevotella | 0.5314 |
| RuminiclostridiumC | CAG-83 | 0.5311 |
| CAG-103 | CAG-110 | 0.5277 |
| CAG-110 | Alistipes | 0.5275 |
| Acetatifactor | Roseburia | 0.5255 |
| Paraprevotella | BacteroidesA | 0.5253 |
| Phil12 | Oscillibacter | 0.5246 |
| RC9 | BacteroidesB | 0.5244 |
| EubacteriumR | RuminococcusD | 0.5238 |
| Acutalibacter | Ruthenibacterium | 0.5234 |
| RC9 | 43-108 | 0.5197 |
| UBA11774 | EubacteriumR | 0.5191 |
| Pauljensenia | Mogibacterium | 0.5182 |
| RuminiclostridiumE | RuminococcusE | 0.5182 |
| RuminiclostridiumE | Adlercreutzia | 0.5172 |
| RC9 | Alistipes | 0.5167 |
| Phil12 | 43-108 | 0.5153 |
| Turicibacter | Intestinibacter | 0.5152 |
| Ruthenibacterium | ClostridiumA | 0.5142 |
| Erysipelatoclostridium | Agathobacter | 0.5141 |
| Paraprevotella | 43-108 | 0.5136 |
| UBA11774 | Lachnospira | 0.5132 |
| Phil12 | Parabacteroides | 0.5115 |
| Gabonibacter | BacteroidesB | 0.5096 |
| CAG-180 | RuminococcusC | 0.5092 |
| Butyricimonas | Parabacteroides | 0.5091 |
| Gemmiger | Fusicatenibacter | 0.5069 |
| ER4 | Alistipes | 0.506 |
| Megasphaera | CAG-127 | 0.5057 |
| CAG-103 | EubacteriumR | 0.5048 |
| UBA11774 | CAG-180 | 0.5022 |
| Barnesiella | BacteroidesB | 0.5018 |
| Pauljensenia | Gordonibacter | 0.5017 |
| UBA11774 | Dialister | 0.5012 |
| Butyricimonas | BacteroidesA | 0.5007 |
| CAG-81 | ClostridiumM | 0.5006 |
| Turicibacter | Romboutsia | 0.5006 |
| Barnesiella | UBA11524 | 0.5 |
| CAG-83 | Alistipes | 0.4999 |
| RuminiclostridiumE | Acidaminococcus | 0.4996 |
| Intestinibacter | Romboutsia | 0.4993 |
| ER4 | CAG-110 | 0.4984 |
| RuminiclostridiumE | CAG-127 | 0.498 |
| Gordonibacter | Ruthenibacterium | 0.497 |
| RuminiclostridiumE | RuminococcusC | 0.4965 |
| Parabacteroides | BacteroidesA | 0.4945 |
| CAG-127 | CAG-180 | 0.4938 |
| Paraprevotella | Parabacteroides | 0.4927 |
| Phil12 | Barnesiella | 0.4923 |
| Collinsella | Bifidobacterium | 0.492 |
| EubacteriumG | Fusicatenibacter | 0.4919 |
| Gabonibacter | Paraprevotella | 0.4905 |
| Acutalibacter | UBA1417 | 0.4869 |
| UBA11774 | TF01-11 | 0.4864 |
| UBA11774 | KLE1615 | 0.4862 |
| RC9 | Barnesiella | 0.486 |
| Gordonibacter | Eggerthella | 0.4851 |

Table S7: Top genus pairs in MS (left) and controls (right).

| Genus 1 | Genus 2 | $\rho_{MS}$ | $\rho_C$ | $\rho_{MS} - \rho_C$ |
|---|---|---|---|---|
| Escherichia | LactobacillusB | 0.4514 | -4e-04 | 0.4519 |
| AlistipesA | Erysipelatoclostridium | 0.2685 | -0.1684 | 0.4368 |
| CAG-1427 | UBA11524 | 0.5895 | 0.1696 | 0.4199 |
| Escherichia | PeH17 | 0.3193 | -0.0969 | 0.4162 |
| UBA1417 | Collinsella | 0.4302 | 0.0311 | 0.3991 |
| EubacteriumF | RuminococcusC | 0.4808 | 0.0898 | 0.391 |
| UBA11774 | RuminiclostridiumC | 0.5192 | 0.1322 | 0.3871 |
| Eggerthella | CAG-180 | 0.3604 | -0.026 | 0.3865 |
| CAG-127 | EubacteriumF | 0.4056 | 0.0386 | 0.367 |
| CAG-177 | Holdemanella | 0.2382 | 0.6056 | -0.3673 |
| Acidaminococcus | CAG-103 | -0.1442 | 0.2238 | -0.368 |
| RC9 | RuminiclostridiumC | -0.106 | 0.262 | -0.368 |
| EubacteriumR | LactobacillusB | -0.1249 | 0.244 | -0.3689 |
| UBA11774 | RuminococcusD | 0.1998 | 0.569 | -0.3691 |
| Escherichia | EubacteriumF | -0.0738 | 0.2961 | -0.3699 |
| EubacteriumI | CAG-56 | 0.1131 | 0.483 | -0.37 |
| Catenibacterium | RuminococcusE | 0.1005 | 0.4707 | -0.3702 |
| UBA11774 | KLE1615 | 0.1154 | 0.4862 | -0.3708 |
| CAG-180 | Alistipes | -0.0924 | 0.2787 | -0.3712 |
| CAG-177 | Dialister | -0.1301 | 0.2425 | -0.3726 |
| Holdemanella | BacteroidesB | -0.2571 | 0.1158 | -0.3729 |
| EubacteriumR | RuminococcusD | 0.1506 | 0.5238 | -0.3732 |
| CAG-180 | Dialister | -0.0913 | 0.2845 | -0.3758 |
| RuminiclostridiumC | Gemmiger | 0.0292 | 0.4051 | -0.3759 |
| PeH17 | 43-108 | 0.0029 | 0.3793 | -0.3764 |
| Sellimonas | CAG-81 | 0.1679 | 0.5457 | -0.3778 |
| PeH17 | Barnesiella | 0.0195 | 0.3987 | -0.3792 |
| Barnesiella | 43-108 | 0.2832 | 0.6627 | -0.3795 |
| Paraprevotella | PeH17 | -0.1372 | 0.2444 | -0.3816 |
| RuminococcusD | LactobacillusB | -0.0761 | 0.3072 | -0.3834 |
| UBA11524 | Prevotella | -0.0192 | 0.3657 | -0.3849 |
| UBA1417 | Alistipes | -0.0484 | 0.3381 | -0.3866 |
| AlistipesA | Catenibacterium | -0.157 | 0.2309 | -0.3879 |
| CAG-180 | Prevotella | -0.036 | 0.3524 | -0.3885 |
| UBA11774 | CAG-180 | 0.1136 | 0.5022 | -0.3886 |
| CAG-103 | BacteroidesB | -0.1354 | 0.2553 | -0.3907 |
| Barnesiella | LactobacillusB | -0.2596 | 0.1336 | -0.3933 |
| Megasphaera | LactobacillusB | 0.1493 | 0.5432 | -0.3939 |
| CAG-1427 | BacteroidesB | -0.172 | 0.2225 | -0.3944 |
| TF01-11 | Lachnospira | 0.1733 | 0.5684 | -0.3951 |
| CAG-83 | BacteroidesB | -0.0479 | 0.349 | -0.397 |
| Romboutsia | Dialister | -0.0883 | 0.3087 | -0.397 |
| EubacteriumI | Dialister | -0.1052 | 0.2931 | -0.3983 |
| CAG-103 | Prevotella | -0.0523 | 0.3489 | -0.4012 |
| Acidaminococcus | ClostridiumA | -0.1407 | 0.2626 | -0.4033 |
| Senegalimassilia | RuminiclostridiumC | -0.1995 | 0.2057 | -0.4052 |
| RC9 | PeH17 | -0.0939 | 0.315 | -0.4089 |
| UBA11524 | BacteroidesB | -0.0873 | 0.3258 | -0.4131 |
| PeH17 | Butyricimonas | -0.1548 | 0.2588 | -0.4136 |
| Paraprevotella | UBA11524 | -0.1174 | 0.2997 | -0.4171 |
| OlsenellaE | Butyricimonas | -0.4047 | 0.0125 | -0.4172 |
| 43-108 | Holdemanella | -0.1889 | 0.2283 | -0.4173 |
| Romboutsia | Lachnospira | -0.1476 | 0.2708 | -0.4184 |
| Acidaminococcus | PeH17 | -0.1147 | 0.3044 | -0.4192 |
| OlsenellaE | Alistipes | -0.3759 | 0.0437 | -0.4196 |
| Dialister | LactobacillusB | -0.2289 | 0.191 | -0.4199 |
| KLE1615 | Lachnospira | 0.2561 | 0.6761 | -0.42 |
| Acidaminococcus | Acutalibacter | -0.0654 | 0.3582 | -0.4236 |
| Holdemanella | Butyricimonas | -0.2845 | 0.1416 | -0.4261 |
| CAG-170 | Alistipes | 0.0214 | 0.4482 | -0.4268 |
| Phil12 | PeH17 | -0.1518 | 0.2756 | -0.4274 |
| Megasphaera | RuminococcusE | -0.0315 | 0.4023 | -0.4338 |
| CAG-127 | Dialister | 0.0493 | 0.4834 | -0.434 |
| Megasphaera | Adlercreutzia | -0.0649 | 0.3716 | -0.4365 |
| Catenibacterium | RuminococcusD | -0.1425 | 0.2963 | -0.4389 |
| UBA11774 | Prevotella | -0.0422 | 0.3981 | -0.4403 |
| Megasphaera | Coprococcus | -0.0857 | 0.3615 | -0.4472 |
| EubacteriumR | Lachnospira | -0.1536 | 0.305 | -0.4586 |
| Megasphaera | CAG-170 | -0.1098 | 0.3512 | -0.461 |
| Acidaminococcus | RuminococcusD | 0.0112 | 0.4729 | -0.4618 |
| CAG-83 | Alistipes | 0.0357 | 0.4999 | -0.4642 |
| CAG-180 | RuminococcusD | 0.082 | 0.5499 | -0.4679 |
| 43-108 | UBA1417 | -0.0965 | 0.375 | -0.4715 |
| Gabonibacter | TF01-11 | -0.0565 | 0.4158 | -0.4723 |
| Acidaminococcus | CAG-170 | -0.1144 | 0.365 | -0.4794 |
| UBA1417 | BacteroidesB | -0.2295 | 0.2525 | -0.4819 |
| EubacteriumR | Prevotella | -0.203 | 0.2838 | -0.4868 |
| Dialister | UBA1417 | -0.071 | 0.4202 | -0.4913 |
| UBA11774 | Acidaminococcus | -0.0573 | 0.4377 | -0.495 |
| RuminiclostridiumE | LactobacillusB | -0.1886 | 0.3084 | -0.4971 |
| Senegalimassilia | Adlercreutzia | -0.2275 | 0.2735 | -0.501 |
| RuminiclostridiumC | CAG-83 | 0.0213 | 0.5311 | -0.5098 |
| PeH17 | BacteroidesB | -0.2355 | 0.278 | -0.5135 |
| Adlercreutzia | Dialister | -0.293 | 0.2326 | -0.5256 |
| RuminiclostridiumE | Acidaminococcus | -0.0328 | 0.4996 | -0.5323 |
| Acidaminococcus | CAG-177 | -0.0945 | 0.4461 | -0.5406 |
| Megasphaera | RuminococcusC | -0.0671 | 0.4807 | -0.5478 |
| Megasphaera | CAG-127 | -0.0442 | 0.5057 | -0.5499 |
| Acidaminococcus | EubacteriumR | -0.2135 | 0.3547 | -0.5683 |
| Megasphaera | CAG-180 | -0.1967 | 0.3853 | -0.5819 |
| Acidaminococcus | Adlercreutzia | -0.1906 | 0.413 | -0.6036 |
| EubacteriumR | Dialister | -0.2587 | 0.3482 | -0.6069 |
| Megasphaera | CAG-177 | -0.1881 | 0.4253 | -0.6135 |
| Megasphaera | RuminococcusD | -0.1457 | 0.4686 | -0.6143 |
| Acidaminococcus | CAG-180 | -0.2094 | 0.4055 | -0.6149 |
| RuminiclostridiumE | Megasphaera | -0.0811 | 0.5342 | -0.6154 |
| UBA11774 | Lachnospira | -0.1046 | 0.5132 | -0.6178 |
| Acidaminococcus | RuminococcusC | -0.0643 | 0.563 | -0.6273 |
| Acidaminococcus | CAG-127 | -0.0385 | 0.6221 | -0.6606 |
| UBA11774 | Dialister | -0.1607 | 0.5012 | -0.6619 |

Table S8: Top 100 differences in MS ($\rho_{MS}$) versus controls ($\rho_C$).

Parimal Mukhopadhyay. *Complex Surveys: Analysis of Categorical Data*. Springer, 2016.

Chuan Tian. Microbial network recovery by compositional graphical Lasso under additive log-ratio transformation. 2020.
